# Efficient estimation for large-scale linkage disequilibrium patterns of the human genome

**DOI:** 10.1101/2023.06.18.545475

**Authors:** Xin Huang, Tian-Neng Zhu, Ying-Chao Liu, Guo-An Qi, Jian-Nan Zhang, Guo-Bo Chen

**Affiliations:** Institute of Bioinformatics, Zhejiang University, Hangzhou, Zhejiang Province, China; Center for General Practice Medicine, Department of General Practice Medicine; Center for Reproductive Medicine, Department of Genetic and Genomic Medicine, and Clinical Research Institute, Zhejiang Provincial People’s Hospital, People’s Hospital of Hangzhou Medical College, Hangzhou, Zhejiang, China; Alibaba Group, Hangzhou, Zhejiang, China; Key Laboratory of Endocrine Gland Diseases of Zhejiang Province, Hangzhou, Zhejiang, China; Hainan Institute of Zhejiang University, Sanya, Hainan, China

## Abstract

In this study, we proposed an efficient algorithm (X-LD) for estimating LD patterns for a genomic grid, which can be of inter-chromosomal scale or of small segments. Compared with conventional methods, the proposed method was significantly faster, dropped from 𝒪 (*nm*^2^) to 𝒪 (*n*^*2*^*m*)—*n* the sample size and *m* the number of SNPs, and consequently we were permitted to explore in depth unknown or reveal long-anticipated LD features of the human genome. Having applied the algorithm for 1000 Genome Project (1KG), we found: **I**) The extended LD, driven by population structure, was universally existed, and the strength of inter-chromosomal LD was about 10% of their respective intra-chromosomal LD in relatively homogeneous cohorts, such as FIN and to nearly 56% in admixed cohort, such as ASW. **II**) After splitting each chromosome into upmost more than a half million grids, we elucidated the LD of the HLA region was nearly 42 folders higher than chromosome 6 in CEU and 11.58 in ASW; on chromosome 11, we observed that the LD of its centromere was nearly 94.05 folders higher than chromosome 11 in YRI and 42.73 in ASW. **III**) We uncovered the long-anticipated inversely proportional linear relationship between the length of a chromosome and the strength of chromosomal LD, and their Pearson’s correlation was on average over 0.80 for 26 1KG cohorts. However, this linear norm was so far perturbed by chromosome 11 given its more completely sequenced centromere region. Uniquely chromosome 8 of ASW was found most deviated from the linear norm than any other autosomes. The proposed algorithm has been realized in C++ (called X-LD) and available at https://github.com/gc5k/gear2, and can be applied to explore LD features in any sequenced populations.

## Introduction

Linkage disequilibrium (LD) is the association for a pair of loci and the metric of LD serves as the basis for developing genetic applications in agriculture, evolutionary biology, and biomedical researches (Weir, 2008; Hill and Robertson, 1966). The structure of LD of the human genome is shaped by many factors, mutation, recombination, population demography, epistatic fitness, and completeness of genomic data itself (Myers *et al*., 2005; Nei and Li, 1973; Ardlie *et al*., 2002). Due to its overwhelming cost, LD structure investigation is often compromised to a small genomic region (Chang *et al*., 2015; Theodoris *et al*., 2021), and their typical LD structure is as illustrated for a small segment (Barrett *et al*., 2005). Now, given the availability of large-scale genomic data, such as millions of single nucleotide polymorphisms (SNPs), the large-scale LD patterns of the human genome play crucial roles in determining genomics studies, and many theories and useful algorithms upon large-scale LD structure, from genome-wide association studies, polygenic risk prediction for complex diseases, and choice for reference panels for genotype imputation (Vilhjálmsson *et al*., 2015; Yang and Zhou, 2020; Bulik-Sullivan *et al*., 2015; Yang *et al*., 2011; Das *et al*., 2016).

However, there are impediments, largely due to intensified computational cost, in both investigating large-scale LD and providing high-resolution illustration for their details. If we consider a genomic grid that is consisted of *m*^*2*^ SNP pairs, given a sample of *n* individuals and *m* SNPs (*n «m)* – typically as observed in 1000 Genomes Project (1KG) (Lowy-Gallego *et al*., 2019), its benchmark computational time cost for calculating all pairwise LD is 𝒪 (*nm*^2^), a burden that quickly drains computational resources given the volume of the genomic data. In practice, it is of interest to know the mean LD of the 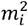 SNP pairs for a genomic grid, which covers *m*_*i*_ *×m*_*j*_ SNP pairs. Upon how a genomic grid is defined, a genomic grid consequently can be consisted of : **i**) the whole genome-wide *m*^*2*^ SNP pairs, and we denote their mean LD as *ℓg* ; **ii**) the intra-chromosomal mean LD for the *ith* chromosome of 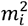 SNP pairs, and denote as *ℓi* ; **iii**) the inter-chromosomal mean LD and chromosomal *m*_*i*_ *m*_*j*_ SNP pairs, and denoted as *ℓi· j* .

In this study we propose an efficient algorithm that can estimate *ℓg, ℓi*, and *ℓ i· j* ·, the computational time of which can be reduced from 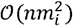 to 𝒪 (*n*^*2*^*m*_*i*_) for *ℓi* and 𝒪 (*nm*_*i*_ *m*_*j*_) to 𝒪 (*n*^*2*^*m*_*i*_*+n*^*2*^*m*_*j*_ ) for *ℓi· j*. The rationale of the proposed method relies on the connection between the genetic relationship matrix (GRM) and LD (Chen, 2014; Goddard, 2009), and in this study a more general transformation from GRM to LD can be established via Isserlis’s theorem (Isserlis, 1918; Zhou, 2017). The statistical properties, such as sampling variance, of the estimated LD have been derived too.

The proposed method can be analogously considered a more powerful realization for Haploview (Barrett *et al*., 2005), but additional utility can be derived to bring out unprecedented survey of LD patterns of the human genome. As demonstrated in 1KG, we consequently investigate how biological factors such as population structure, admixture, or variable local recombination rates can shape large-scale LD patterns of the human genomes.

1. The proposed method provides statistically unbiased estimates for large-scale LD patterns and shows computational merits compared with the conventional methods **(Figure 2)**.
2. We estimated *ℓg*, and 22 autosomal *ℓi*. and 231 inter-autosomal *ℓi· j* for the 1KG cohorts. There were ubiquitously existence of extended LD, which was associated with population structure or admixture **(Figure 3)**.
3. We provided high-resolution illustration that decomposed a chromosome into upmost nearly a million grids, each of which was consisted of 250×250 SNP pairs, the highest resolution that has been realized so far at autosomal level **(Figure 4)**; tremendous variable recombination rates led to regional strong LD as highlighted for the HLA region of chromosomes 6 and the centromere region of chromosome 11.
4. Furthermore, a consequently linear regression constructed could quantify LD decay score genome-widely, and in contrast LD decay was previously surrogated in a computational expensive method. There was strong ethnicity effect that was associated with extended LD **(Figure 5)**.
5. We demonstrate that the strength of autosomal *ℓi*. was inversely proportional to the SNP number, an anticipated relationship that is consistent to genome-wide spread of recombination hotspots. However, the chromosome 8 of ASW showed substantial deviation from the fitted linear relationship **(Figure 6)**.

**Figure 1.**
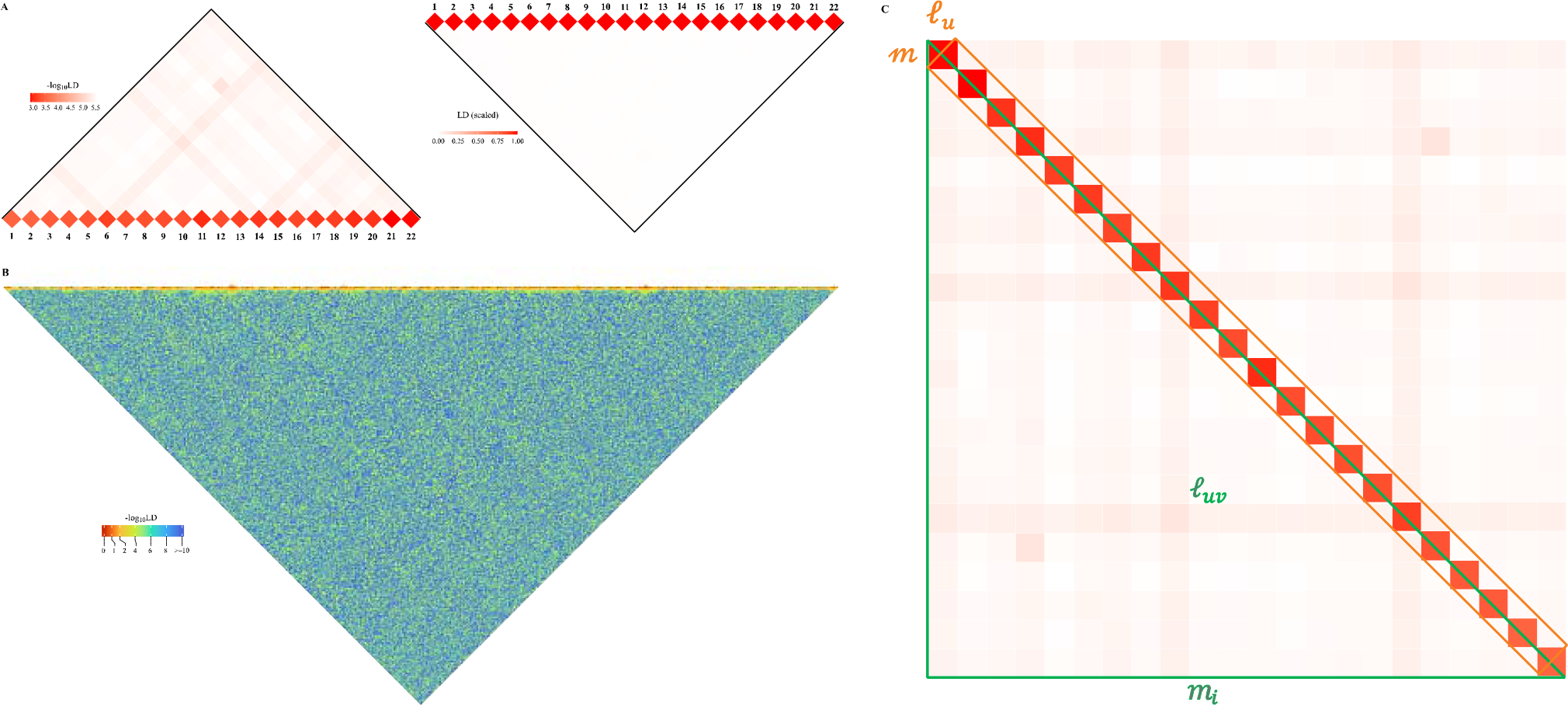
Schematic illustration for large-scale LD analysis as exampled for CONVERGE cohort. **A)** The 22 human autosomes have consequently 22 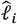 and 231 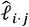, ut (left) and with (right) scaling transformation; Scaling transformation is given in **Eq 8. B**) If zoom into chromosome 2 of 420,946 SNPs, a chromosome of relative lity is expected to have self-similarity structure that harbors many approximately strong 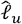 along the diagonal, and relatively weak 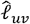 off-diagonally. Here osome 2 of CONVERGE has been split into 1,000 blocks and yielded 1000 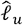 LD grids, and 499,500 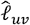 LD grids. **C**) An illustration of the construction process for D-decay regression model

**Figure 2.**
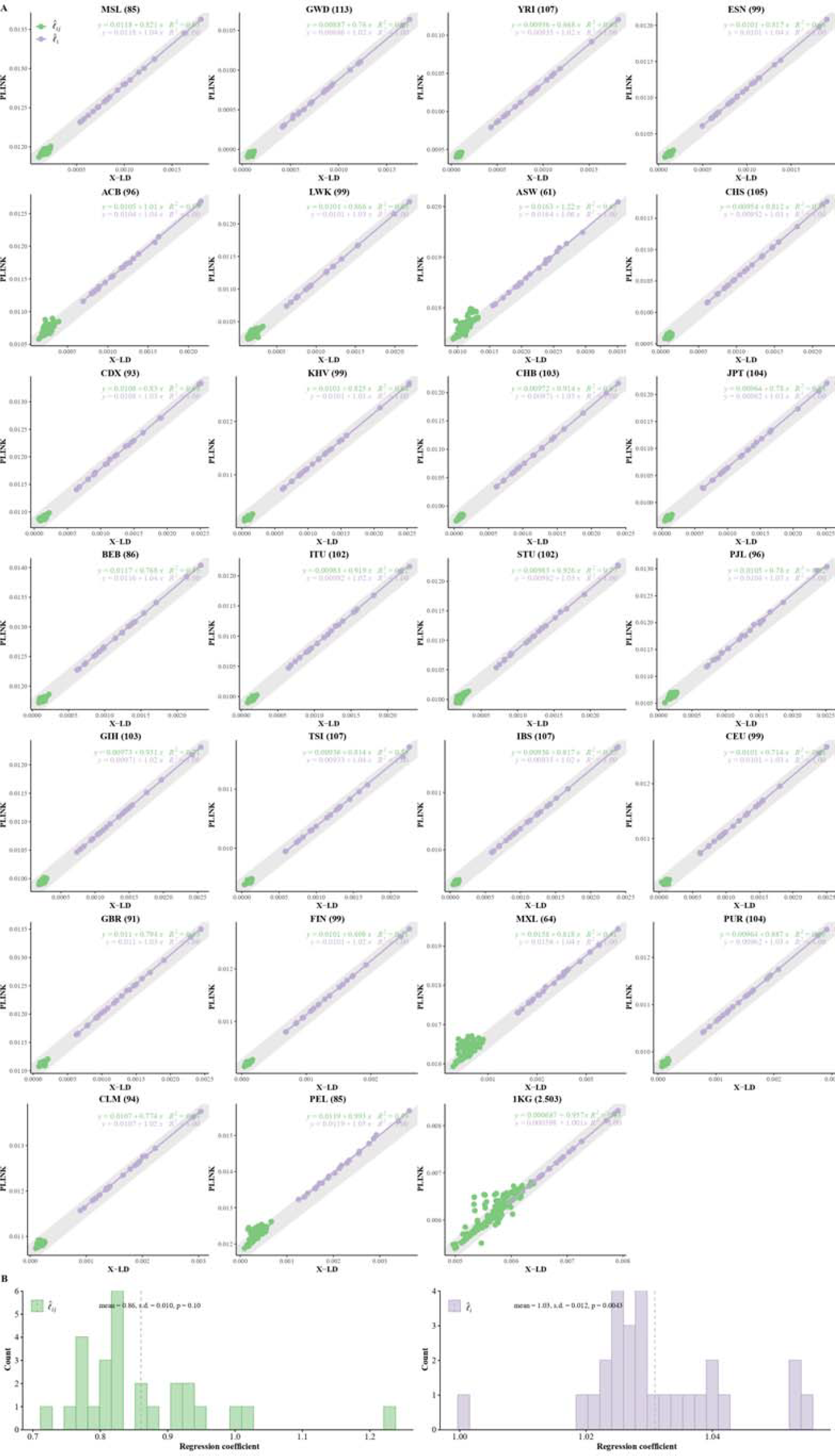
Reconciliation for LD estimators in the 26 cohorts of 1KG. **A)** Consistency examination for the 26 1KG cohorts for their 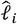 and 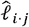 estimated by X-LD and PLINK (--r2). In each figure, the 22 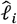 fitting line is in purple, whereas the 231 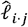 fitting line is in green. The gray solid line,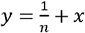, in which *n* the sample size of each cohort, represents the expected fit between PLINK and X-LD estimates, and the two estimated regression models at the top-right corner of each plot shown this consistency. The sample size of each cohort is in parentheses. **B)** Distribution of *R*^2^ of 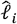 and 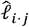 fitting lines is based on X-LD and PLINK algorithms in the 26 cohorts; *R*^2^ represents variation explained by the fitted model. 26 1KG cohorts: MSL (Mende in Sierra Leone), GWD (Gambian in Western Division, The Gambia), YRI (Yoruba in Ibadan, Nigeria), ESN (Esan in Nigeria), ACB (African Caribbean in Barbados), LWK (Luhya in Webuye, Kenya), ASW (African Ancestry in Southwest US); CHS (Han Chinese South), CDX (Chinese Dai in Xishuangbanna, China), KHV (Kinh in Ho Chi Minh City, Vietnam), CHB (Han Chinese in Beijing, China), JPT (Japanese in Tokyo, Japan); BEB (Bengali in Bangladesh), ITU (Indian Telugu in the UK), STU (Sri Lankan Tamil in the UK), PJL (Punjabi in Lahore, Pakistan), GIH (Gujarati Indian in Houston, TX); TSI (Toscani in Italia), IBS (Iberian populations in Spain), CEU (Utah residents (CEPH) with Northern and Western European ancestry), GBR (British in England and Scotland), FIN (Finnish in Finland); MXL (Mexican Ancestry in Los Angeles, California), PUR (Puerto Rican in Puerto Rico), CLM (Colombian in Medellin, Colombia), PEL (Peruvian in Lima, Peru).

**Figure 3.**
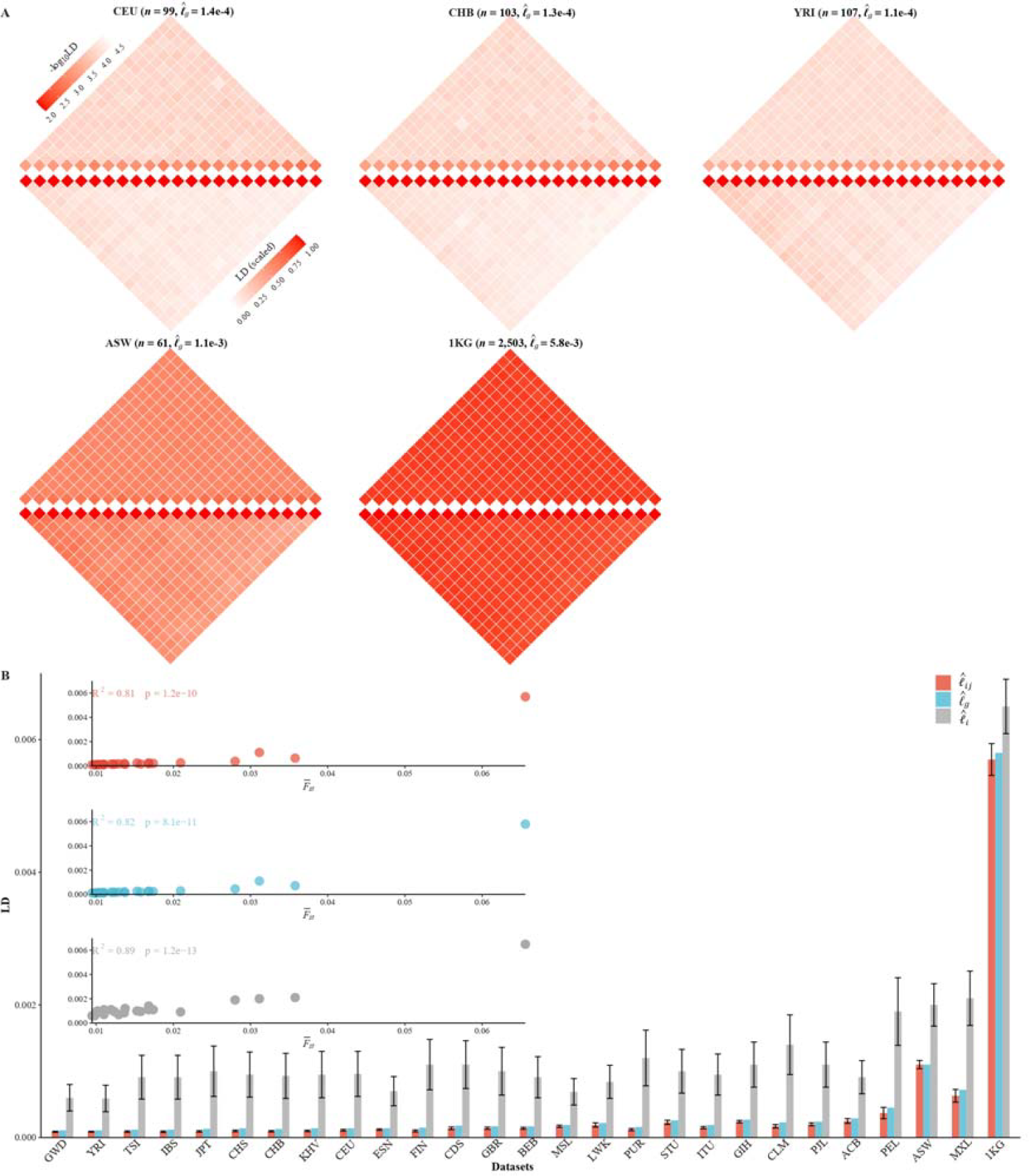
Various LD components for the 26 1KG cohorts. **A)** Chromosomal scale LD components for 5 representative cohorts (CEU, CHB, YRI, ASW, and 1KG). The upper parts of each figure represent 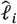 (along the diagonal) and 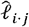 (off-diagonal), and the lower part 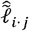 as in **Eq 8**. For visualization purposes, the quantity of LD before scaling is transformed to a -log10 scale, with smaller values (red hues) representing larger LD, and a value of 0 representing that all SNPs are in LD. **B)** The relationship between the degree of population structure (approximated by 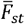) and 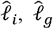, and 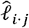 in the 26 1KG cohorts.

**Figure 4.**
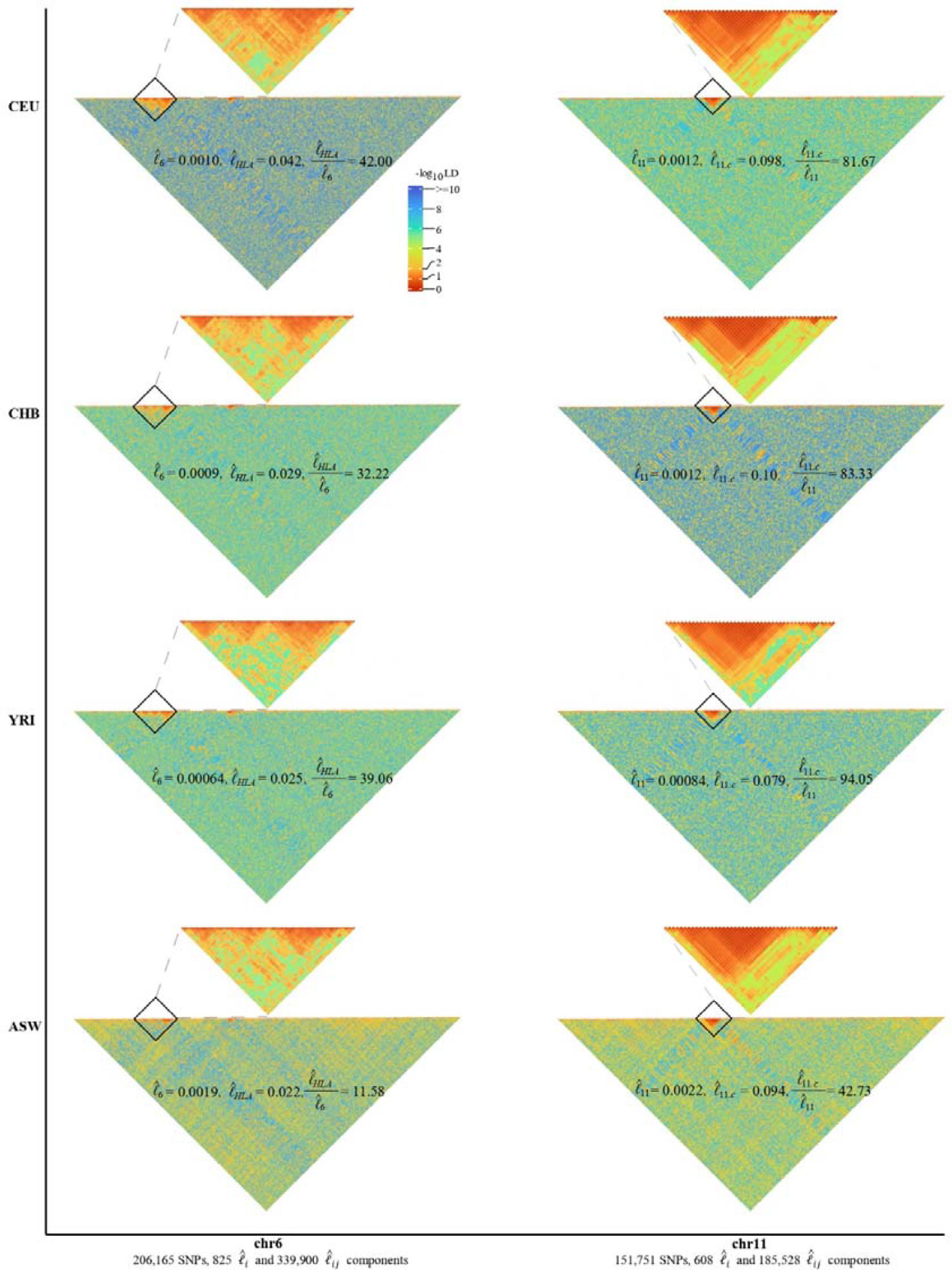
High-resolution illustration for LD grids for CEU, CHB, YRI, and ASW (m 250). For each cohort, we partition chromosomes 6 and 11 into high-resolution LD grids (each LD grid contains 250 X 250 SNP pairs). The bottom half of each figure shows the LD grids for the entire chromosome. Further zooming into HLA on chromosome 6 and the centromere region on chromosome 11, and their detailed LD in the relevant regions are also provided in the upper half of each figure. For visualization purposes, LD is transformed to a -log10-scale, with smaller values (red hues) representing larger LD, and a value of 0 representing that all SNPs are in LD.

**Figure 5.**
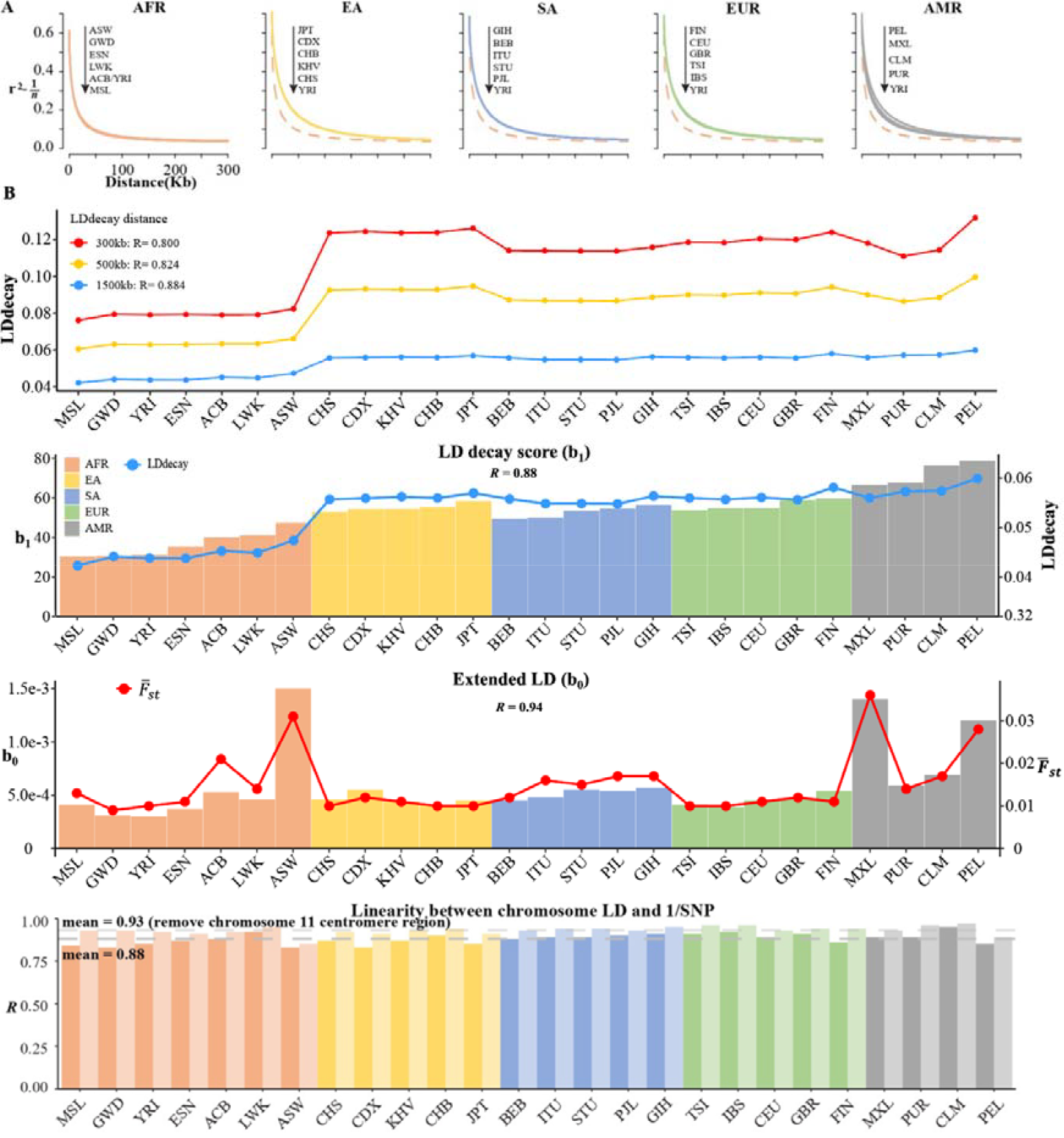
LD decay analysis for 26 1KG cohorts. **A)** Conventional LD decay analysis in PLINK for 26 cohorts. To eliminate the influence of sample size, the inverse of sample size has been subtracted from the original LD values. The YRI cohort, represented by the orange dotted line, is chosen as the reference cohort in each plot. The top-down arrow shows the order of LDdecay values according to **Table 5. B)** Model-based LD decay analysis for the 26 1KG cohorts. We regressed each autosomal against its corresponding inversion of the SNP number for each cohort. Regression coefficient quantifies the averaged LD decay of the genome and intercept provides a direct estimate of possible existence of long-distance LD. The values in the first three plot indicate the correlation between and LD decay score in three different physical distance and the correlation between 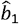 (left-side vertical axis) and LD decay score (right-side vertical axis) and the correlation between 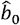 (left-side vertical axis) and 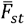 (right-side vertical axis), respectively. The last plot assessed the impact of centromere region of chromosome 11 on the linear relationship between chromosomal LD and the inverse of the SNP number. The dark and light gray dashed lines represent the mean of the ℛwith and without the presence of centromere region of chromosome 11.

**Figure 6.**
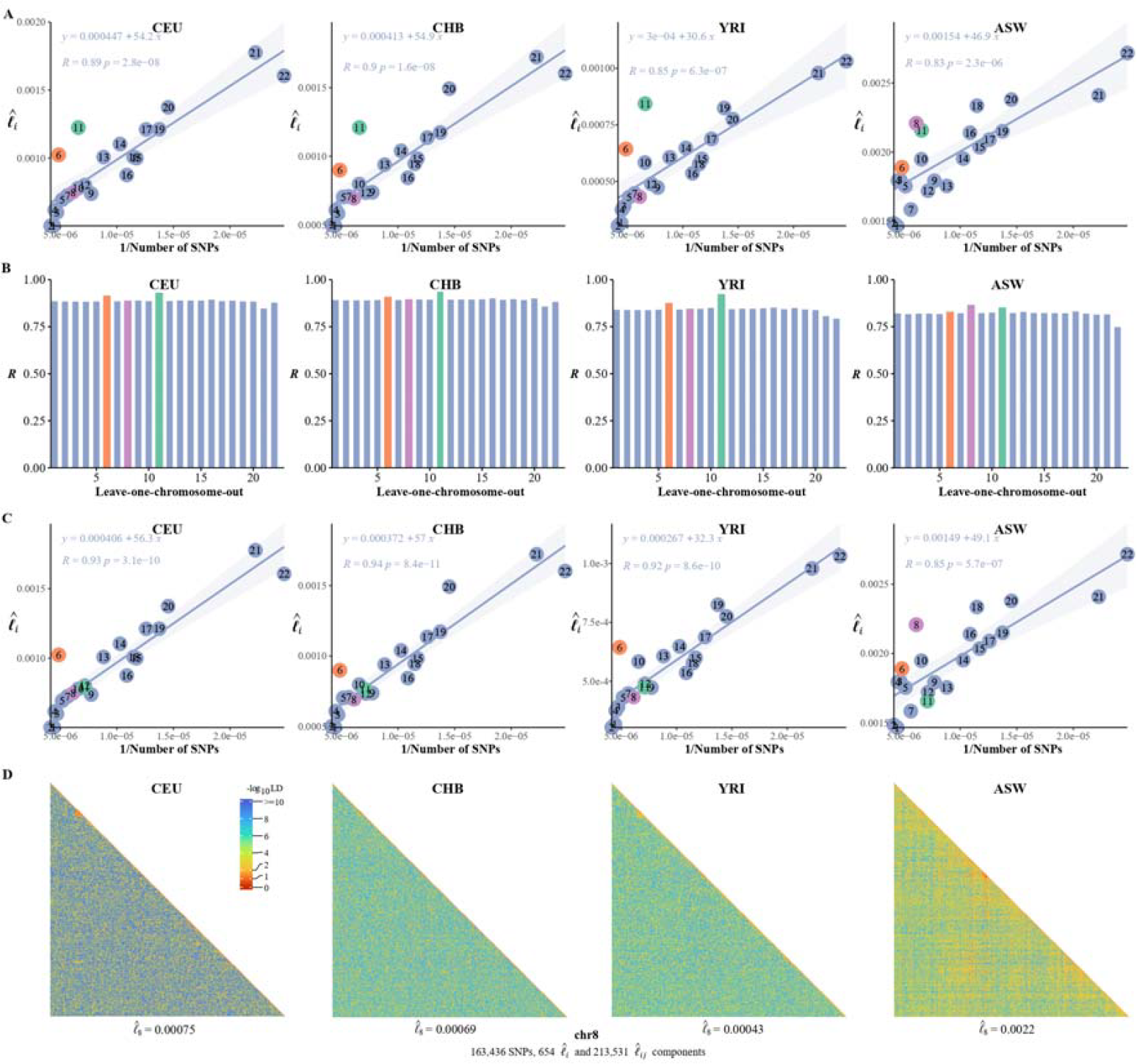
The correlation between the inversion of the SNP number and 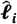. **A)** The correlation between the inversion of the SNP number and 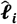 in CEU, CHB, YRI, and ASW. **B)** Leave-one-chromosome-out strategy is adopted to evaluate the contribution of a certain chromosome on the correlation between the inverse of the SNP number and 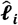 **C)** The correlation between the inversion of the SNP number and chromosomal LD in CEU, CHB, YRI, and ASW after removing the centromere region of chromosome 11. **D)** High-resolution illustration for LD grids for chromosome 8 in CEU, CHB, YRI, and ASW. For each cohort, we partition chromosome 8 into consecutive LD grids (each LD grid contains 250 X 250 SNP pairs). For visualization purposes, LD is transformed to a -log10-scale, with smaller values (red hues) representing larger LD, and a value of 0 representing that all SNPs are in LD.

The proposed algorithm has been realized in C++ and is available at: https://github.com/gc5k/gear2. As tested the software could handle sample size as large as 10,000 individuals.

## Methods and Materials

### The overall rationale for large-scale LD analysis

We assume LD for a pair of biallelic loci is measured by squared Pearson’s correlation, 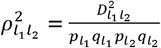 in which 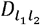 the LD of loci *l*_1_ and *l*_2_, *p* and *q* the reference and the alternative allele frequencies. If we consider the averaged LD for a genomic grid over 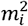 SNP pairs, the conventional estimator is 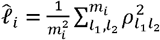 and, if we consider the averaged LD for *m*_*i*_ *and m*_*j*_ SNP pairs between two genomic segments, then 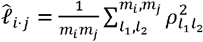 . Now let us consider the 22 human autosomes (**Figure 1A**). We naturally partition the genome into 𝒞=22 blocks, and its genomic LD, denoted as *ℓg* can be expressed as

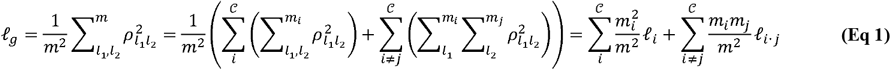

So we can decompose *ℓg* into 𝒞 *ℓi*. and 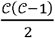 unique *ℓi*· *j* . Obviously, **Eq 1** can be also expressed in the context for a single chromosome 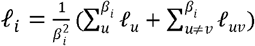, in which 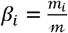 the number of SNP segments, each of which has *m* SNPs. Geometrically it leads to β_*i*_ diagonal grids and 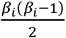 unique off-diagonal grids **(Figure 1B)**.

### LD-decay regression

As human genome can be boiled down to small LD blocks by genome-widely spread recombination hotspots (Hinch *et al*., 2019; Li *et al*., 2022), mechanically there is self-similarity for each chromosome that the relatively strong *ℓi*. for juxtaposed grids along the diagonal but weak *ℓi· j* for grids slightly off-diagonal. So, for a chromosomal *ℓi*., we can further express it as

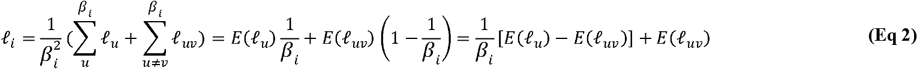

in which *ℓu* is the mean LD for a diagonal grid, *ℓuv* the mean LD for off-diagonal grids, and *m*_*i*_ the number of SNPs on the *i*^th^ chromosome. Consider a linear model below (**Figure 1C** for its illustration),

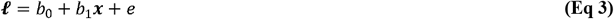

in which ***ℓ*** represents a vector composed of 𝒞 *ℓi*., ***x*** represents a vector composed of 𝒞 *x*_*i*_ and 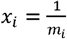 the inversion of the SNP number of the chromosome. The regression coefficient and intercept can be estimated as below

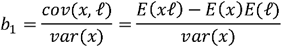

And

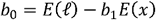

There are some technical details in order to find the interpretation for *b*_0_ and *b*_1_ .We itemize them briefly. For the mean and variance of *x*:

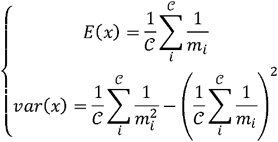

For *E*(*xℓ)*

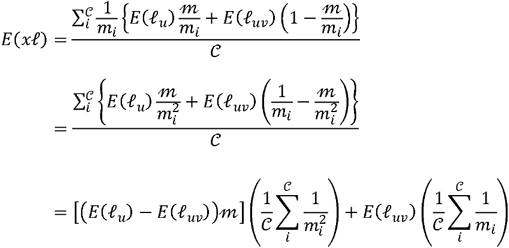

For *E*(*x)* For *E*(*ℓ)*

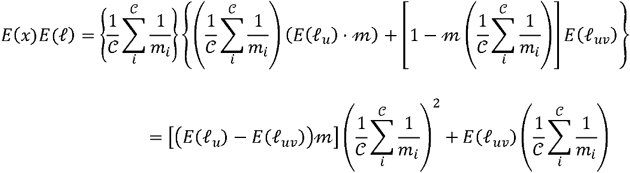

Then we integrate these items to have the expectation for *b*_1_:

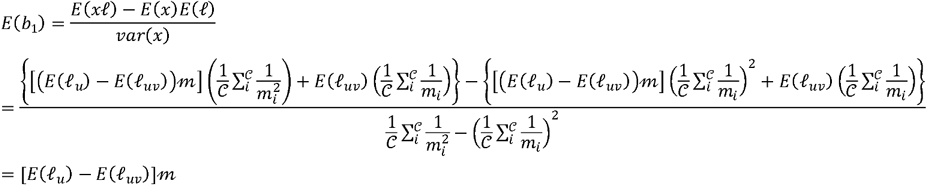

d

Similarly, we plug in *E*(*b*_1_) so as to derive *b*_0_ :

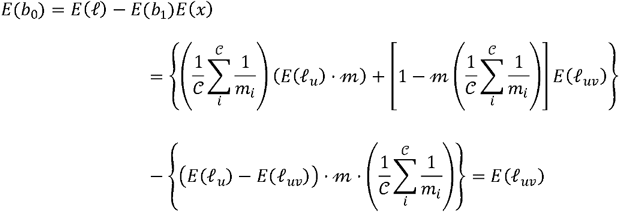

After some algebra, if *E*(*ℓ*_*u*_) *≫ E* (*ℓ*_*uv*_) – say if the former is far greater than the latter, the interpretation of *b*_1_ and *b*_0_ can be

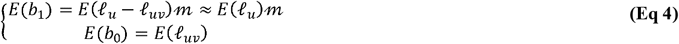

It should be noticed that *E*(*b*_*1*_) ≈ *E*(*ℓ*_*u*_)*m* quantifies the averaged LD decay of the genome. Conventional LD decay is analysed via the well-known LD decay analysis, but **Eq 4** provides a direct estimate of both LD decay and possible existence of extended LD. We will see the application of the model in **Figure 5** that the strength of the long-distance LD is associated with population structure. of note, the underlying assumption of **Eq 3** and **Eq 4** is genome-wide spread of recombination hotspots, an established result that has been revealed and confirmed (Hinch *et al*., 2019).

### Efficient estimation for ℓ_*g*_, ℓ_*i*_, and ℓ_*i ·j*_

For the aforementioned analyses, the bottleneck obviously lies in the computational cost in estimating *ℓ*_*i*_ and *ℓ*_*i·j·*_ *ℓ*_*i*_ and *ℓ*_*i·j*_ are used to be estimated via the current benchmark algorithm as implemented in PLINK (Chang *et al*., 2015), and the computational time complex is proportional to *O*(*nm*^*2*^). We present a novel approach to estimate *ℓ*_*i*_ and *ℓ*_*i·j·*_ Given a genotypic matrix X, a *n × m* mmatrix, if we assume that there are *m*_*i*_ and *m*_*j*_ SNPs on chromosomes *i* and *j*, respectively, we can construct *n × n* genetic relatedness matrices as below

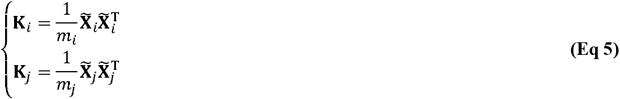

in which 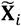 is the standardized ***X***_*i*_ and 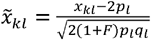 where *x*_*kl*_ is the genotype for the *K*^*th*^ individual at the *l*^*th*^ biallelic locus, *F* is the inbreeding coefficient having the value of 0 for random mating population and 1 for an inbred population, *p*_*l*_ and *q*_*l*_ are the frequencies of the reference and the alternative alleles (*p*_*l*_ + *q*_*l*_ =1)respectively. When GRM is given, we can obtain some statistical characters of ***K***_*i*_. We extract two vectors 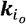, which stacks the off-diagonal elements of ***K***_*i*_, and 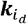 which takes the diagonal elements of ***K***_*i*_. The mathematical expectation of 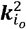 in which 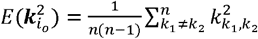 can be established according to Isserlis’s theorem in terms of the four-order moment (Isserlis, 1918),

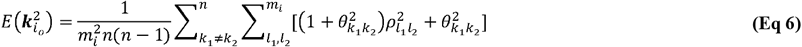

in which 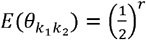 is the expected relatedness score and *r* indicates the *r*^*th*^ degree relatives. *r*=0 for the same individual, and *r*=1 for the first degree relatives. Similarly, we can derive for 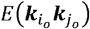 **Eq 6** establishes the connection between GRM and the aggregated LD estimation that 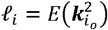 .According to Delta method as exampled in Appendix I of Lynch and Walsh’s book (Lynch and Walsh, 1998), the means and the sampling variances for *ℓ*_*i*_ and *ℓ*_*i·j*_ are,

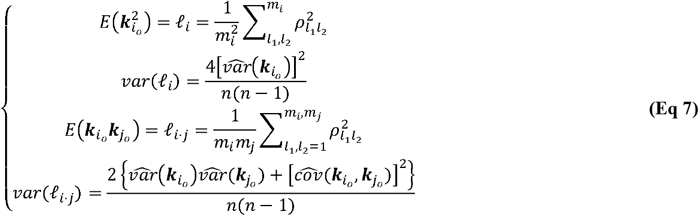

in which 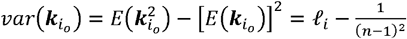 and 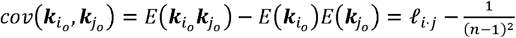, respectively. Of note, the properties of *ℓ*_*g*_ can be derived similarly if we replace *ℓ*_*i*_ with *ℓ*_*g*_ in **Eq 7**. We can develop 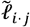, a scaled version of *ℓ*_*i·j·*_ as below

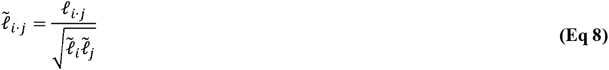

in which 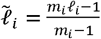, a modification that removed the LD with itself. According to Delta method, the sampling variance of 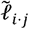 is

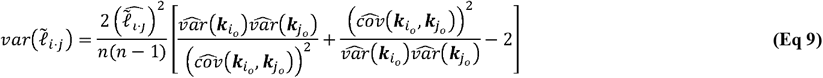

Of note, when there is no LD between a pair of loci, *ℓ* yields zero and its counterpart PLINK estimate yields 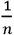,a difference that can be reconciled in practice **(see Figure 2)**.

### Raise of LD due to population structure

In this study, the connection between LD and population structure is bridged via two pathways below, in terms of a pair of loci and of the aggregated LD for all pair of loci. For a pair of loci, their LD is often simplified as 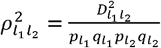, but will be inflated if there are subgroups (Nei and Li, 1973). In addition, it is well established the connection between population structure and eigenvalues, and in particular the largest eigenvalue is associated with divergence of subgroups (Patterson *et al*., 2006). In this study, the existence of subgroups of cohort is surrogated by the largest eigenvalue 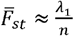.

### Data description and quality control

The 1KG (Auton *et al*., 2015), which is launched to produce a deep catalogue of human genomic variation by whole genome sequencing (WGS) or whole exome sequencing (WES), and 2,503 strategically selected individuals of global diversity are included (containing 26 cohorts). We used the following criteria for SNP inclusion for each of the 26 1KG cohorts: i) autosomal SNPs only; ii) SNPs with missing genotype rates higher than 0.2 were removed, and missing genotypes were imputed; iii) Only SNPs with minor allele frequencies higher than 0·05 were retained. Then 2,997,635 consensus SNPs that were present in each of the 26 cohorts were retained. According to their origins, the 26 cohorts are grouped as African (AFR: MSL, GWD, YRI, ESN, ACB, LWK, and ASW), European (EUR: TSI, IBS, CEU, GBR, and FIN), East Asian (EA: CHS, CDX, KHV, CHB, and JPT), South Asian (SA: BEB, ITU, STU, PJL, and GIH), and American (AMR: MXL, PUR, CLM, and PEL), respectively.

In addition, to test the capacity of the developed software (X-LD), we also included CONVERGE cohort (*n*=10,640), which was used to investigate Major Depressive Disorder (MDD) in the Han Chinese population (Cai *et al*., 2015). We performed the same criteria for SNP inclusion as that of the 1KG cohorts, and *m=*5,215,820 SNPs were remained for analyses.

### X-LD software implementation

The proposed algorithm has been realized in our X-LD software, which is written in C++ and reads in binary genotype data as often used in PLINK. As multi-thread programming is adopted, the efficiency of X-LD can be improved upon the availability of computational resources. We have tested X-LD in various independent datasets for its reliability and robustness. Certain data management options, such as flexible inclusion or exclusion of chromosomes, have been built into the commands of X-LD. In X-LD, missing genotypes are naively imputed according to Hardy-Weinberg proportions; however, when missing rate is high, we suggest the genotype matrix should be imputed by other advanced imputation tools.

The most time-consuming part of X-LD was the construction of GRM 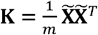, and the established computational time complex was *𝒪* (*n*^2^*m*). However, if 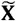 is decomposed into 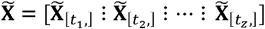, in which 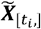, has dimension of *n* × *B*, using Mailman algorithm the computational time complex for building **K** can be reduced to 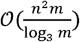 (Liberty and Zucker, 2009). This idea of embedding Mailman algorithm into certain high throughput genomic studies has been successful, and our X-LD software is also leveraged by absorbing its recent practice in genetic application (Wu and Sankararaman, 2018).

## Results

### Statistical properties of the proposed method

**Table 1** introduced symbols frequently cited in this study. As schematically illustrated in **Figure 1**, *ℓ*_*g*_ could be decomposed into *𝒞 ℓ*_*i*_ and 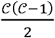 unique *ℓ*_*i,j*_ components. We compared the estimated *ℓ;*_*i*_ and *ℓ*_*i,j*_ in X-LD with those being estimated in PLINK (known as “--r2”, and the estimated squared Pearson’s correlation LD is denoted as *r*^2^). Considering the 212 substantial computational cost of PLINK, only 100,000 randomly selected autosome SNPs were used for each 1KG cohort, and 22 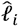 and 231 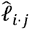 were estimated. After regressing 22 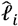 against those of PLINK, we found that the regression slope was close to unity and bore an anticipated intercept a quantity of approximately 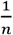 **(Figure 2A and Figure 2B)**.

**Table 1.** Notation definitions.

| Notation | Definition |
| --- | --- |
| $\mathcal{C}$ | The number of chromosomes. |
| $i$ and $j$ | Subscripts index chromosome $i$ and $j$. |
| $\beta_i$ | The number of SNP segments of chromosome $i$, each of which has $m$ SNPs. |
| $D_{l_1 l_2}$ | The difference between the observed and expected haplotype frequencies, with $D_{l_1 l_2} = p_{l_1 l_2} - p_{l_1} p_{l_2}$. |
| $F$ | The inbreeding coefficient. |
| $\mathbf{K}_i$ | Genetic relatedness matrix for chromosome $i$, and two vectors, $\mathbf{k}_{i_o}$ and $\mathbf{k}_{i_d}$, from $\mathbf{K}_i$, where $\mathbf{k}_{i_o}$ stacks the off-diagonal elements and $\mathbf{k}_{i_d}$ stacks the diagonal elements. |
| $k$ | Subscript indexes individual. |
| $l_1$ and $l_2$ | Subscripts index a pair of SNPs. |
| $m$ | The number of SNPs; $m_i$ the number of SNPs on chromosome $i$. |
| $n$ | The number of samples; $n_i$, the number of samples in subpopulation $i$. |
| $p_l$ and $q_l$ | Frequency of the $l^{th}$ reference allele and alternative allele in the population. |
| $\theta_{k_1 k_2}$ | The relatedness score between individual $k_1$ and $k_2$. |
| $x_{kl}$ | The genotype for the $k^{th}$ individual at the $l^{th}$ biallelic locus. |
| $\mathbf{X}_i$ and $\tilde{\mathbf{X}}_i$ | Genotype and standardized genotype matrixes for chromosome $i$. |
| $\rho_{l_1 l_2}^2$ | Squared Pearson's correlation coefficient for any pair of SNPs, including a SNP to itself when $l_1 = l_2$. |
| $r^2$ | Squared Pearson's correlation metric for LD but estimated from PLINK (--r2) or PopLDdecay. |
| $\ell_g$ | The mean LD of the whole genome-wide $m^2$ SNP pairs. |
| $\ell_i$ | The intra-chromosomal mean LD for the $i^{th}$ chromosome of $m_i^2$ SNP pairs. |
| $\ell_{i,j}$ | The inter-chromosomal mean LD $i^{th}$ and $j^{th}$ chromosomal $m_i m_j$ SNP pairs, a scaled version is $\tilde{\ell}_{ij}$. |
| $\ell_u$ | The mean LD for a diagonal grid. |
| $\ell_{uv}$ | The mean LD for off-diagonal grids. |

In other words, PLINK gave 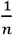 even for SNPs of no LD. However, when regressing 231 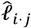 estimates against those of PLINK, it was found that largely because of tiny quantity of 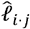 it was slightly smaller than 1 but statistically insignificant from 1 in these 26 1KG cohorts (mean of 0.86 and s.d. of 0.10, and its 95 % confidence interval was (0.664, 1.056)); when the entire 1KG samples were used, its much larger LD due to subgroups, nearly no estimation bias was found **(Figure 2A and Figure 2B)**. In contrast, because of their much larger values, 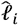 components were always consistent with their corresponding estimates from PLINK (mean of 1.03 and s.d. of 0.012, 95% confidence interval was (1.006, 1.053), bearing an ignorable bias). Furthermore, we also combined the African cohorts together (MSL, GWD, YRI, ESN, LWK, totaling 599 individuals), the East Asian cohorts together (CHS, CDX, KHV, CHB, and JPT, totaling 504 individuals), and the European cohorts together (TSI, IBS, CEU, GBR, and FIN, totaling 503 individuals), the resemblance pattern between X-LD and PLINK was similar as observed in each cohort alone **(Figure S1)**. The empirical data in 1KG verified that the proposed method was sufficiently accurate.

To fairly evaluate the computational efficiency of the proposed method, the benchmark comparison was conducted on the first chromosome of the entire 1KG dataset (*n* =2.503) and *m* =(225,967), and 10 CPUs were used for multi-thread computing. Compared with PLINK, the calculation efficiency of X-LD was nearly 30∼40 times faster for the tested chromosome, and its computational time of X-LD was proportional to 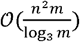 **(Figure S2)**. So, X-LD provided a feasible and reliable estimation of large-scale complex LD patterns. More detailed computational time of the tested tasks would be reported in their corresponding sections below; since each 1KG cohort had sample size around 100, otherwise specified the computational time was reported for CHB (*n* = 103) as a reference **(Table 2)**. In order to test the capability of the software, the largest dataset tested was CONVERGE (*n* =10,640, and (*n* =5,215,820), and it took 77,508.00 seconds, about 22 hours, to estimate 22 autosomal 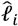 and 231 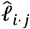 **(Figure 1A)**;

**Table 2.** Computational time for the demonstrated estimation tasks.

| Cohort | Task description | Time cost | Computational time complex |
| --- | --- | --- | --- |
| CHB ($n = 103$, $m = 2,997,655$) | Estimation for 22 autosomal $\ell_i$, and 231 inter-chromosomal $\ell_{i,j}$. Results see **Figure 3 and Table 3.** | 101,34 secs | $\mathcal{O}(n^2m)$ |
| 1KG ($n = 2,503$, $m = 2,997,655$) | Same as above. | 3,008.29 secs | Same as above |
| CONVERGE ($n = 10,640$, $m = 5,215,820$) | Same as above. Result see **Figure 1A.** | 77,508.00 secs | Same as above |
| **Estimation for high-resolution LD interaction given bin size of 250 SNPs** |  |  |  |
| CHB ($n = 103$, $m_2 = 241,241$) | Chromosome 2, estimation for 965 $\ell_i$, and 465,130 $\ell_{i,j}$. Results see **Figure 4.** | 66.86 secs | $\mathcal{O}\left(n^2\left(m_i + \left(\frac{m_i}{250}\right)^2\right)\right)$ |
| CHB ($n = 103$, $m_{22} = 40,378$) | Chromosome 22, estimation for 162 $\ell_i$, and 13,041 $\ell_{i,j}$. Results see **Figure 4.** | 3.22 secs | Same as above |
| CONVERGE ($n = 10,640$, $m_{22} = 71,407$) | Chromosome 22, estimation for 286 $\ell_i$, and 40,755 $\ell_{i,j}$. | 8,736.29 secs | Same as above |
| CONVERGE ($n = 10,640$, $m_2 = 420,949$) | Chromosome 2, estimation for 1,000 $\ell_i$, and 499,500 $\ell_{i,j}$. Result see **Figure 1B.** | 45,125.00 secs | Chromosome 2 was split into 1000 blocks, each of which had about 420 SNPs. |

**Notes:** for the sake of fair comparison, 10 CPUs were used for multi-thread computing.

When zooming into chromosome 2 of CONVERGE, on which 420,949 SNP had been evenly split into 1,000 blocks and yielded 1,000 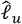 grids, and 499,500 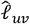 LD grids, it took 45,125.00 seconds, about 12.6 hours, to finished the task **(Figure 1B)**.

### Ubiquitously extended LD and population structure/admixture

We partitioned the 2,997,635 SNPs into 22 autosomes **(Figure 3A and Figure S3)**, and the general LD patterns were as illustrated for CEU, CHB, YRI, ASW, and 1KG. As expected, 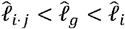 for each cohort **(Figure 3B)**. As observed in these 1KG cohorts, all these three LD measures were associated with population structure, which was surrogated by 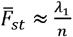, and their squared correlation *R*^2^ were greater than 0.8. ACB, ASW, PEL, and MXL, which all showed certain admixture, tended to have much greater 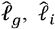, and 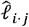 **(Table 3 and Figure 3B)**. In contrast, East Asian (EA) and European (EUR) orientated cohorts, which showed little within cohort genetic differentiation – as their largest eigenvalues were slightly greater than 1, had their aggregated LD relatively low and resembled each other **(Table 3)**. Furthermore, for several European (TSI, IBS, and FIN) and East Asian (JPT) cohorts, the ratio between 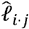 and 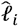 components could be smaller than 0.1, and the smallest ratio was found to be about 0.091 in FIN. The largest ratio was found in 1KG that 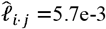 and 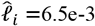, and the ratio was 0.877 because of the inflated LD due to population structure. A more concise statistic to describe the ratio between *ℓ*_*i,j*_and *ℓ;*_*i*_ was 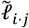, **Eq 8**, and the corresponding values for 231 scaled 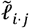 for FIN was 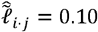 (s.d. of 0.027) and for 1KG was 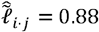 (s.d. of 0.028).

**Table 3.** X-LD estimation for complex LD components (2,997,635 SNPs)

| Cohort (n) | Ancestry | $\lambda_1(\bar{F}_{st})^1$ | $\hat{\ell}_g$ (s.e.)<sup>2</sup> | $\hat{\ell}_i$ (s.d.)<sup>3</sup> | $\hat{\ell}_{i,j}$ (s.d.)<sup>3</sup> | $\hat{\ell}_{i,j}$ (s.d.)<sup>3</sup> | Lower bound of LD<sup>4</sup> |
| --- | --- | --- | --- | --- | --- | --- | --- |
| MSL (85) | AFR | 1.10 (0.013) | 1.9e-4 (1.21e-6) | 6.9e-4 (2.0e-4) | 1.7e-4 (1.7e-5) | 0.26 (0.053) | 0.161971831 |
| GWD (113) | AFR | 1.07 (0.009) | 1.1e-4 (5.61e-7) | 6.0e-4 (2.0e-4) | 8.7e-5 (8.1e-6) | 0.16 (0.037) | 0.247218789 |
| YRI (107) | AFR | 1.05 (0.010) | 1.1e-4 (4.23e-7) | 5.9e-4 (2.0e-4) | 8.8e-5 (6.9e-6) | 0.16 (0.04) | 0.242001641 |
| ESN (99) | AFR | 1.09 (0.011) | 1.4e-4 (7.67e-7) | 7.0e-4 (2.2e-4) | 1.2e-4 (1.2e-5) | 0.19 (0.043) | 0.217391304 |
| ACB (96) | AFR | 2.01 (0.021) | 2.9e-4 (3.78e-6) | 9.1e-4 (2.5e-4) | 2.5e-4 (3.6e-5) | 0.29 (0.070) | 0.147727273 |
| LWK (99) | AFR | 1.35 (0.014) | 2.2e-4 (2.38e-6) | 8.4e-4 (2.5e-4) | 1.9e-4 (3.2e-5) | 0.24 (0.052) | 0.173913043 |
| ASW (61) | AFR | 1.90 (0.031) | 1.1e-3 (2.73e-5) | 2.0e-3 (3.2e-4) | 1.1e-3 (6.2e-5) | 0.57 (0.059) | 0.079681275 |
| CHS (105) | EA | 1.08 (0.010) | 1.4e-4 (9.39e-7) | 9.5e-4 (3.4e-4) | 1.0e-4 (1.3e-5) | 0.12 (0.030) | 0.31147541 |
| CDX (93) | EA | 1.11 (0.012) | 1.8e-4 (1.38e-6) | 1.1e-3 (3.6e-4) | 1.4e-4 (2.0e-5) | 0.14 (0.040) | 0.272277228 |
| KHV (99) | EA | 1.07 (0.011) | 1.4e-4 (7.67e-7) | 9.5e-4 (3.5e-4) | 1.0e-4 (1.2e-5) | 0.12 (0.031) | 0.31147541 |
| CHB (103) | EA | 1.07 (0.010) | 1.3e-4 (6.94e-7) | 9.3e-4 (3.4e-4) | 9.5e-5 (1.1e-5) | 0.11 (0.030) | 0.317948718 |
| JPT (104) | EA | 1.06 (0.010) | 1.3e-4 (7.22e-7) | 1.0e-3 (3.8e-4) | 9.3e-5 (1.2e-5) | 0.10 (0.028) | 0.338638673 |
| BEB (86) | SA | 1.07 (0.012) | 1.7e-4 (8.09e-7) | 9.1e-4 (3.1e-4) | 1.4e-4 (1.5e-5) | 0.17 (0.042) | 0.236363636 |
| ITU (102) | SA | 1.61 (0.016) | 1.9e-4 (1.84e-6) | 9.5e-4 (3.1e-4) | 1.5e-4 (1.7e-5) | 0.18 (0.044) | 0.231707317 |
| STU (102) | SA | 1.56 (0.015) | 2.6e-4 (3.21e-6) | 1.0e-3 (3.3e-4) | 2.3e-4 (3.1e-5) | 0.23 (0.047) | 0.171526587 |
| PJL (96) | SA | 1.67 (0.017) | 2.4e-4 (2.74e-6) | 1.1e-3 (3.4e-4) | 2.0e-4 (2.2e-5) | 0.21 (0.048) | 0.20754717 |
| GIH (103) | SA | 1.73 (0.017) | 2.7e-4 (3.41e-6) | 1.1e-3 (3.4e-4) | 2.4e-4 (1.9e-5) | 0.23 (0.049) | 0.179153094 |
| TSI (107) | EUR | 1.07 (0.010) | 1.2e-4 (6.10e-7) | 9.1e-4 (3.3e-4) | 9.0e-5 (1.1e-5) | 0.11 (0.029) | 0.325 |
| IBS (107) | EUR | 1.07 (0.010) | 1.2e-4 (6.10e-7) | 9.1e-4 (3.3e-4) | 8.8e-5 (1.1e-5) | 0.11 (0.028) | 0.329949239 |
| CEU (99) | EUR | 1.07 (0.011) | 1.4e-4 (7.67e-7) | 9.6e-4 (3.4e-4) | 1.1e-4 (1.3e-5) | 0.12 (0.030) | 0.293577982 |
| GBR (91) | EUR | 1.11 (0.012) | 1.7e-4 (1.08e-6) | 1.0e-3 (3.6e-4) | 1.4e-4 (1.8e-5) | 0.15 (0.036) | 0.253807107 |
| FIN (99) | EUR | 1.09 (0.011) | 1.5e-4 (9.69e-7) | 1.1e-3 (3.8e-4) | 1.0e-4 (1.5e-5) | 0.10 (0.027) | 0.34375 |
| MXL (64) | AMR | 2.29 (0.036) | 7.2e-4 (1.49e-5) | 2.1e-3 (4.1e-4) | 6.3e-4 (9.6e-5) | 0.32 (0.072) | 0.136986301 |
| PUR (104) | AMR | 1.43 (0.014) | 1.6e-4 (1.30e-6) | 1.2e-3 (4.2e-4) | 1.2e-4 (1.7e-5) | 0.11 (0.026) | 0.322580645 |
| CLM (94) | AMR | 1.58 (0.017) | 2.3e-4 (2.49e-6) | 1.4e-3 (4.5e-4) | 1.7e-4 (2.6e-5) | 0.13 (0.035) | 0.281690141 |
| PEL (85) | AMR | 2.38 (0.028) | 4.5e-4 (7.33e-6) | 1.9e-3 (5.1e-4) | 3.7e-4 (8.5e-5) | 0.21 (0.062) | 0.196483971 |
| 1KG (2,503) | MIX | 164.20 (0.066) | 5.8e-3 (4.63e-6) | 6.5e-3 (4.1e-4) | 5.7e-3 (2.4e-4) | 0.88 (0.028) | 0.051505547 |

<sup>1</sup>Eigenvalue was estimated. In parentheses was the ratio between the listed largest eigenvalue and the sample size. Since it exists an approximation that $\bar{F}_{st} \approx \frac{\lambda_1}{n}$ , the ratio can be taken as an approximation of population structure.
<sup>2</sup> Standard error was calculated as $\frac{2}{\sqrt{n(n-1)}}[\hat{\ell}_g - \frac{1}{(n-1)^2}]$ , as **Eq 7**.
<sup>3</sup> Estimated empirically from $\mathcal{C}$ chromosomal $\hat{\ell}_i$ ; Estimated empirically from $\frac{\mathcal{C}(\mathcal{C}-1)}{2}$ inter-chromosomal $\hat{\ell}_{i,j}$ .
<sup>4</sup> It is estimated by $\frac{22\hat{\ell}_i}{22\hat{\ell}_i + 231\hat{\ell}_{i,j}}$ , indicating lower bound of true LD.

In terms of computational time, for 103 CHB samples, it took about 101.34 seconds to estimate 22 autosomal 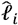 and 231 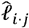 ; for all 1KG 2,503 samples, X-LD took about 3,008.29 seconds **(Table 1)**. Conventional methods took too long to complete the analyses in this section, so no comparable computational time was provided. For detailed 22 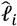 and 231 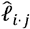 estimates for each 1KG cohort, please refer to **Extended Data 1 (Excel Sheet 1-27)**.

### Detecting exceedingly high LD grids shaped by variable recombination rates

We further explored each autosome with high-resolution grid LD visualization. We set *m* =250, so each grid had the *ℓ;*_*uv*_ for 250 × 250 SNP pairs. The computational time complex was 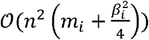, in which 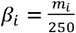, and with our proposed method in CHB it costed 66.86 seconds for chromosome 2, which had 241,241 SNPs and was totaled 466,095 unique grids, and 3.22 seconds only for chromosome 22, which had 40,378 SNPs and was totaled 13,203 unique grids **(Table 1)**. In contrast, under conventional methods those LD grids were not very likely to be exhaustively surveyed because of its computational cost was 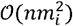: for CHB chromosome 2, it would have taken about 40 hours as estimated. As the result was very similar for *m* =500 **(Figure S4)**, we only reported the results under *m* =250 below.

As expected, chromosome 6 (206,165 SNPs, totaling 340,725 unique grids) had its HLA cluster showing much higher LD than the rest of chromosome 6. In addition, we found very dramatic variation of the HLA cluster LD 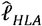 HLA (28,477,797-33,448,354 bp, totaling 3,160 unique grids) across ethnicities. For CEU, CHB, YRI, and ASW, their 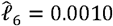, and 0.0019, respectively, but their corresponding HLA cluster grids had 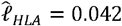 and 0.022, respectively **(Figure 4)**.

Consequently, the largest ratio for 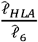 was of 42.00 in CEU, 39.06 in YRI, and 32.22 in CHB, but was reduced to 11.58 in ASW. Before the release of CHM13 (Hoyt *et al*., 2022), chromosome 11 had the most completely sequenced centromere region, which had much rarer recombination events, all four cohorts showed an strong LD 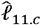 around the centromere (46,061,947-59,413,484 bp, totaling 1,035 unique grids) regardless of their ethnicities **(Figure 4)**. 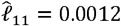, and 0.0022, respectively, and 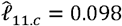, and 0.094, respectively; the ratio for 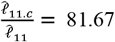, and 94,05, for CEU, CHB, and YRI, respectively; the lowest ratio was found in ASW of 42.73. In addition, removing the HLA region of chromosome 6 or the centromere region of chromosome 11 would significantly reduce 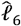 or 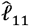 in comparison with the randomly removal of other regions **(Figure S5)**.

### Model-based LD decay regression revealed LD composition

The real LD block size was not exact of *m* =250 or *m* =500, but an unknown parameter that should be inferred in computational intensive “LD decay” analysis (Zhang *et al*., 2019; Chang *et al*., 2015). We conducted the conventional LD decay for the 26 1KG cohorts **(Figure 5A)**, and the time cost was 1,491.94 seconds for CHB. For each cohort, we took the area under the LD decay curve in the LD decay plot, and it quantified approximately the LD decay score for each cohort. The smallest score was 0.0421 for MSL and the largest was 0.0598 for PEL (**Table 5**). However, this estimation was not taken into account the real extent of LD, so it was not precise enough to reflect the LD decay score. For example, for admixture population, such as American cohorts, the extent of LD would be longer.

In contrast, we proposed a model-based method, as given in **Eq 3**, which could estimate LD decay score (regression coefficient *b*_1_) and long-distance LD score (intercept *b*_0_) jointly. Given the estimated 22 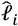 **(Extended data 1; Table 4 for four representative cohorts and Supplementary R code)**, we regressed each autosomal 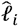 against its correspondingly inversion of SNP number, and all yielded positive slopes (Pearson’s correlation ℛ > 0.80, **Table 5; Figure 5B)**, an observation that was consistent with genome-wide spread of recombination hotspots. This linear relationship could consequently be considered the norm for a relative homogenous population as observed in most 1KG cohorts **(Figure S6)**, while for the all 2,503 1KG samples ℛ =0.55 only **(Table 5)**, indicating that the population structure and possible differentiated recombination hotspots across ethnicities disturbed the assumption underlying **Eq 3** and smeared the linearity. We extracted 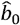 and 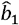 for the 26 1KG cohorts for further analysis. The rates of LD decay score, as indicated by 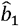, within the African cohorts (AFR) were significantly faster than other continents, consistent with previous observation that African population had relative shorter LD (Gabriel *et al*., 2002); while subgroups within the American continent (AMR) tended to have extended LD range due to their admixed genetic composition **(Table 5** and **Figure 5B)**. Notably, the correlation between 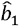 and the approximated LD decay score was ℛ > 0.88. The estimated 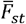 were highly correlated with 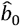.

**Table 4.** Estimates for 22 autosomal 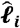 in CEU, CHB, YRI, and ASW, respectively.

| Chromosome | SNP number | $\hat{\ell}_i$ | | | |
| --- | --- | --- | --- | --- | --- |
|  |  | CEU | CHB | YRI | ASW |
| 1 | 225,967 | 5.0e-4 (8.2e-6) | 0.00049 (7.8e-6) | 0.00032 (4.3e-6) | 0.0015 (4e-05) |
| 2 | 241,241 | 5.0e-4 (8.1e-6) | 5.0e-4 (7.9e-6) | 3.0e-4 (4.1e-6) | 0.0015 (4e-05) |
| 3 | 212,670 | 6.0e-04 (1.0e-5) | 0.00058 (9.5e-6) | 0.00039 (5.7e-6) | 0.0018 (5.1e-5) |
| 4 | 222,241 | 0.00062 (1.0e-5) | 0.00061 (1.0e-5) | 0.00038 (5.4e-6) | 0.0018 (5.0e-5) |
| 5 | 193,632 | 0.00069 (1.2e-5) | 7.0e-04 (1.2e-5) | 0.00043 (6.5e-6) | 0.0018 (4.9e-5) |
| 6 | 206,165 | 0.0010 (1.9e-5) | 9.0e-04 (1.6e-5) | 0.00064 (1.0e-5) | 0.0019 (5.4e-5) |
| 7 | 177,414 | 0.00073 (1.3e-5) | 0.00071 (1.2e-5) | 0.00045 (6.8e-6) | 0.0016 (4.3e-5) |
| 8 | 163,436 | 0.00075 (1.3e-5) | 0.00069 (1.2e-5) | 0.00043 (6.5e-6) | 0.0022 (6.4e-5) |
| 9 | 129,440 | 0.00074 (1.3e-5) | 0.00074 (1.3e-5) | 0.00047 (7.2e-6) | 0.0018 (5.0e-5) |
| 10 | 152,251 | 0.00078 (1.4e-5) | 8.0e-04 (1.4e-5) | 0.00058 (9.3e-6) | 0.0019 (5.6e-5) |
| 11 | 151,751 | 0.0012 (2.3e-5) | 0.0012 (2.2e-5) | 0.00084 (1.4e-5) | 0.0022 (6.2e-5) |
| 12 | 139,684 | 8.0e-4 (1.4e-5) | 0.00073 (1.2e-5) | 0.00049 (7.5e-6) | 0.0017 (4.8e-5) |
| 13 | 113,390 | 0.0010 (1.8e-5) | 0.00094 (1.6e-5) | 0.00061 (9.8e-6) | 0.0018 (4.9e-5) |
| 14 | 97,335 | 0.0011 (2.0e-5) | 0.0010 (1.8e-5) | 0.00065 (1.1e-5) | 0.0020 (5.6e-5) |
| 15 | 85,307 | 0.0010 (1.8e-5) | 0.00098 (1.7e-5) | 6.0e-4 (9.6e-6) | 0.0020 (5.8e-5) |
| 16 | 92,007 | 0.00088 (1.6e-5) | 0.00084 (1.5e-5) | 0.00054 (8.4e-6) | 0.0021 (6.2e-5) |
| 17 | 79,478 | 0.0012 (2.3e-5) | 0.0011 (2.0e-5) | 0.00069 (1.1e-5) | 0.0021 (6.0e-5) |
| 18 | 87,105 | 0.0010 (1.8e-5) | 0.00095 (1.7e-5) | 0.00058 (9.2e-6) | 0.0023 (6.8e-5) |
| 19 | 72,794 | 0.0012 (2.3e-05) | 0.0012 (2.1e-5) | 0.00082 (1.4e-5) | 0.0022 (6.2e-5) |
| 20 | 68,881 | 0.0014 (2.6e-5) | 0.0015 (2.7e-5) | 0.00078 (1.3e-5) | 0.0024 (7.0e-5) |
| 21 | 45,068 | 0.0018 (3.4e-5) | 0.0017 (3.2e-5) | 0.00098 (1.7e-5) | 0.0024 (7.1e-5) |
| 22 | 40,378 | 0.0016 (3.1e-5) | 0.0016 (2.9e-5) | 0.0010 (1.8e-5) | 0.0027 (8.1e-5) |

**Note:** each $\hat{\ell}_i$ and its standard error in parentheses, as estimated in Eq 7.

**Table 5.** LD decay regression analysis for 26 cohorts.

| Cohort (n) | LD-decay regression<sup>1</sup> |  |  | Population parameters<sup>2</sup> |  |  |  |
| --- | --- | --- | --- | --- | --- | --- | --- |
| | $\hat{b}_0$ | $\hat{b}_1$ | R | LD decay score | $\bar{F}_{st}$ (%) | Ancestry | True LD<sup>3</sup> |
| MSL (85) | 0.00041 | 29.97 | 0.84 | 0.0421 | 0.013 | AFR | 0.62727273 |
| GWD (113) | 0.00031 | 30.17 | 0.83 | 0.0439 | 0.009 | AFR | 0.65934066 |
| YRI (107) | 0.00030 | 30.64 | 0.85 | 0.0436 | 0.010 | AFR | 0.66292135 |
| ESN (99) | 0.00037 | 34.82 | 0.87 | 0.0436 | 0.011 | AFR | 0.65420561 |
| ACB (96) | 0.00053 | 39.62 | 0.88 | 0.0451 | 0.021 | AFR | 0.63194444 |
| LWK (99) | 0.00046 | 40.52 | 0.92 | 0.0447 | 0.014 | AFR | 0.64615385 |
| ASW (61) | 0.0015 | 46.88 | 0.83 | 0.0472 | 0.031 | AFR | 0.57142857 |
| CHS (105) | 0.00046 | 52.36 | 0.87 | 0.0555 | 0.010 | EA | 0.67375887 |
| CDX (93) | 0.00055 | 53.77 | 0.83 | 0.0557 | 0.012 | EA | 0.66666667 |
| KHV (99) | 0.00044 | 53.79 | 0.87 | 0.0560 | 0.011 | EA | 0.68345324 |
| CHB (103) | 0.00041 | 54.90 | 0.90 | 0.0558 | 0.010 | EA | 0.69402985 |
| JPT (104) | 0.00045 | 57.75 | 0.85 | 0.0568 | 0.010 | EA | 0.68965517 |
| BEB (86) | 0.00045 | 48.84 | 0.88 | 0.0556 | 0.012 | SA | 0.66911765 |
| ITU (102) | 0.00048 | 49.58 | 0.89 | 0.0546 | 0.016 | SA | 0.66433566 |
| STU (102) | 0.00055 | 52.84 | 0.89 | 0.0546 | 0.015 | SA | 0.64516129 |
| PJL (96) | 0.00054 | 54.00 | 0.90 | 0.0546 | 0.017 | SA | 0.67073171 |
| GIH (103) | 0.00057 | 55.81 | 0.91 | 0.0562 | 0.017 | SA | 0.65868263 |
| TSI (107) | 0.00041 | 53.17 | 0.91 | 0.0558 | 0.010 | EUR | 0.68939394 |
| IBS (107) | 0.00039 | 54.22 | 0.92 | 0.0555 | 0.010 | EUR | 0.7 |
| CEU (99) | 0.00045 | 54.23 | 0.89 | 0.0559 | 0.011 | EUR | 0.68085106 |
| GBR (91) | 0.00047 | 58.23 | 0.91 | 0.0555 | 0.012 | EUR | 0.68027211 |
| FIN (99) | 0.00054 | 59.24 | 0.86 | 0.0579 | 0.011 | EUR | 0.67073171 |
| MXL (64) | 0.0014 | 66.13 | 0.89 | 0.0558 | 0.036 | AMR | 0.6 |
| PUR (104) | 0.00059 | 67.20 | 0.89 | 0.0571 | 0.014 | AMR | 0.67039106 |
| CLM (94) | 0.00069 | 75.97 | 0.95 | 0.0572 | 0.017 | AMR | 0.66985646 |
| PEL (85) | 0.0012 | 78.15 | 0.85 | 0.0598 | 0.028 | AMR | 0.61290323 |
| 1KG (2,503) | 0.0061 | 40.65 | 0.55 |  | 0.066 | Mixed | 0.51587302 |

<sup>1</sup>The regression intercept $\hat{b}_0$ and the coefficients $\hat{b}_1$ is as represented in **Eq 3**.
<sup>2</sup>The column for LD decay score was taken the mean of the estimated $r^2 - \frac{1}{n}$ from PopLDdecay in a physical distance of 1500kb, which was approximated to the area under the curve in **Figure 5A** for each cohort; $F_{st}$ was approximated by $\frac{\lambda_1}{n}$ , in which $\lambda_1$ the largest eigenvalue for the cohort. $r^2$ was the estimated LD statistic from PLINK (--r2).
<sup>3</sup>True LD is defined as $\frac{\hat{\ell}_{i,j}}{\hat{\ell}_{i,j} + b_0}$ .

A common feature was universally relative high LD of chromosome 6 and 11 in the 26 1KG cohorts **(Figure S6)**. We quantified the impact of chromosome 6 and 11 by leave-one-chromosome-out test in CEU, CHB, YRI, and ASW for details **(Figure 6A** and **6B)**, and found that dropping chromosome 6 off could lift ℛ on average by 0.017, and chromosome 11 by 0.046. One possible explanation was that the centromere regions of chromosomes 6 and 11 have been assembled more completely than other chromosomes before the completion of CHM13 (Hoyt *et al*., 2022), whereas meiotic recombination tended to be reduced around the centromeres (Hinch *et al*., 2019). We estimated *ℓ;*_*i*_ after having knocked out the centromere region (46,061,947-59,413,484 bp, chr 11) in CEU, CHB, YRI, and ASW, and chromosome 11 then did not deviate much from their respective fitted lines **(Figure 6C)**. A notable exceptional pattern was found in ASW, the chromosome 8 of which had even more deviation than chromosome 11 (ℛ was 0.83 and 0.87 with and without chromosome 8 in leave-one-chromosome out test) **(Figure 6B)**. The deviation of chromosome 8 of ASW was consistent even more SNPs were added **(Figure S7)**. We also provided high-resolution LD grids illustration for chromosome 8 (163,436 SNPs, totaling 214,185 grids) of the four representative cohorts for more detailed virtualization **(Figure 6D)**. ASW had 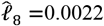, but 0.00075, 0.00069 and 0.00043 for CEU, CHB, and YRI, respectively.

## Discussion

In this study, we present a computationally efficient method to estimate mean LD of genomic grids of many SNP pairs. Our LD analysis framework is based on GRM, which has been embedded in variance component analysis for complex traits and genomic selection (Goddard, 2009; Visscher *et al*., 2014; Chen, 2014). The key connection from GRM to LD is bridged via the transformation between *n* × *n* matrix and *m* × *m* matrix, in particular here via Isserlis’s theorem under the fourth-order moment (Isserlis, 1918). With this connection, the computational cost for estimating the mean LD of *m* × *m* SNP pairs is reduced from 𝒪(*nm*^2^) to 𝒪(*n*^2^*m*), and the statistical properties of the proposed method are derived in theory and validated in 1KG datasets. In addition, as the genotype matrix **X** is of limited entries {0, 1, 2}, assuming missing genotypes are imputed first, using Mailman algorithm the computational cost of GRM can be further reduced to 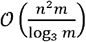 (Liberty and Zucker, 2009). The largest data tested so far for the proposed method has the sample size of 10,640 and of more than 5 million SNPs, it can complete genomic LD analysis in 77,508.00 seconds **(Table 1)**. The weakness of the proposed method is obvious that the algorithm remains slow when the sample size is large or the grid resolution is increased. With the availability of such as UK Biobank data (Bycroft *et al*., 2018), the proposed method may not be adequate, and much advanced methods, such as randomizated implementation for the proposed methods, are needed.

We also applied the proposed method into 1KG and revealed certain characteristics of the human genomes. Firstly, we found the ubiquitously existence of extended LD, which was likely emerged because of population structure, even very slightly, and admixture history. We quantified the 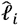 and 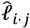 in 1KG, and as indicated by 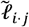 we found the inter-chromosomal LD was nearly an order lower than intra-chromosomal LD; for admixed cohorts, the ratio was much higher, even very close to each other such as in all 1KG samples. Secondly, variable recombination rates shaped peak of local LD. For example, the HLA region showed high LD in European and East Asian cohorts, but relatively low LD in such as YRI, consistent with their much longer population history. Thirdly, it existed general linear correlation between *ℓ;*_*i*_ and the inversion of the SNP number, a long-anticipated result that is as predicted with genome-wide spread of recombination hotspots (Hinch *et al*., 2019). One outlier of this linear norm was chromosome 11, which had so far most completely genotyped centromere and consequently had more elevated LD compared with other autosomes. We anticipate that with the release of CHM13 the linear correlation should be much closer to unity (Hoyt *et al*., 2022). Of note, under the variance component analysis for complex traits, it is often a positive correlation between the length of a chromosome (as surrogated by the number of SNPs) and the proportion of heritability explained (Chen *et al*., 2014).

In contrast, throughout the study recurrent outstanding observations were found in ASW. For example, in ASW the ratio of 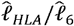 was substantially dropped down compared with that of CEU, CHB, or YRI as illustrated in **Figure 4**. Furthermore, chromosome 8 in ASW fluctuated upwards most from the linear correlation (**Figure 6**), and even after various analyses, such as expanding SNP numbers. One possible explanation may lay under the complex demographic history of ASW, which can be investigated and tested in additional African American samples or possible existence for epistatic fitness (Ni *et al*., 2020).

## Supporting information

ExtendData1

R code for Figure 6C

## Data availability

Public genetic datasets used in this study can be freely downloaded from the following URLs.

**1000 Genomes Project:** https://wwwdev.ebi.ac.uk/eva/?eva-study=PRJEB30460

**CONVERGE:** https://wwwdev.ebi.ac.uk/eva/?eva-study=PRJNA289433

## Acknowledgements

GBC conceived and initiated the study. XH, TNZ, and YL conducted simulation and analyzed data. GBC, XH, TNZ, GAQ, and JZ developed the software. GBC wrote the first draft of the paper. All authors contributed to the writing, discussion of the paper, and validation of the results. This work was supported by National Natural Science Foundation of China (31771392), 110202101032(JY-09), and GZY-ZJ-KJ-23001. The funders played no role in designing, preparation, and submission of the paper.

---

**Figure S1.**
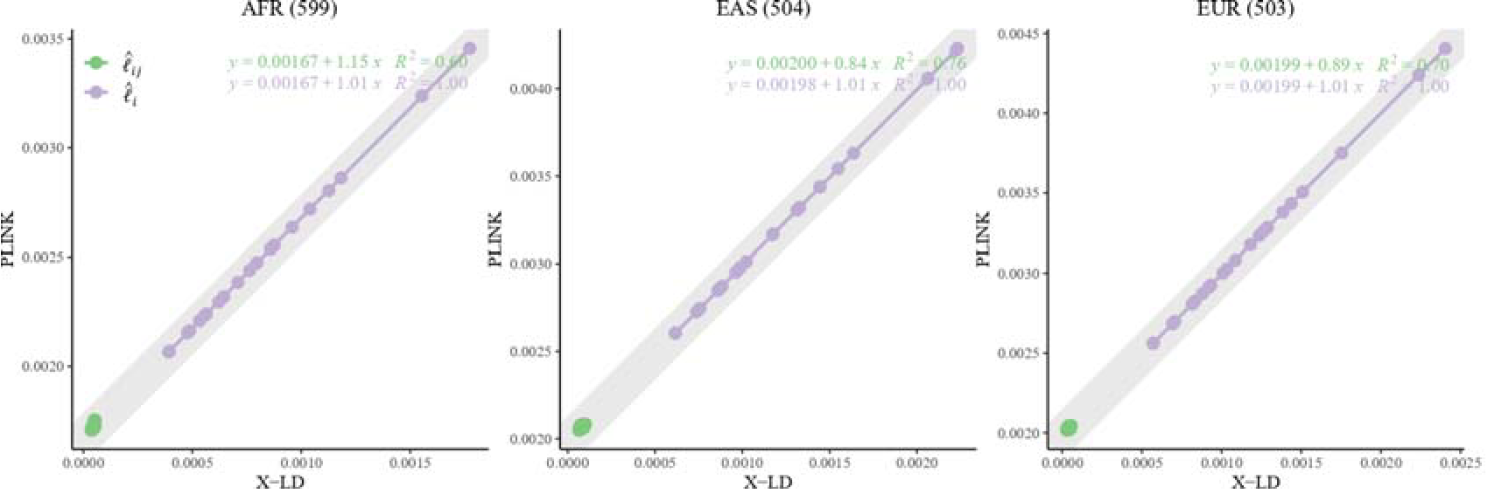
Reconciliation for LD estimators in AFR, EAS, and EUR. In each figure, the 22 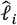 fit line is in purple, whereas the 231 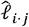 fit line is in green. The gray solid line, 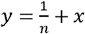, in which *n* the sample size, represents the expected fit between PLINK and X-LD, and the two estimated regression models at the top-right corner shown this consistency.

**Figure S2.**
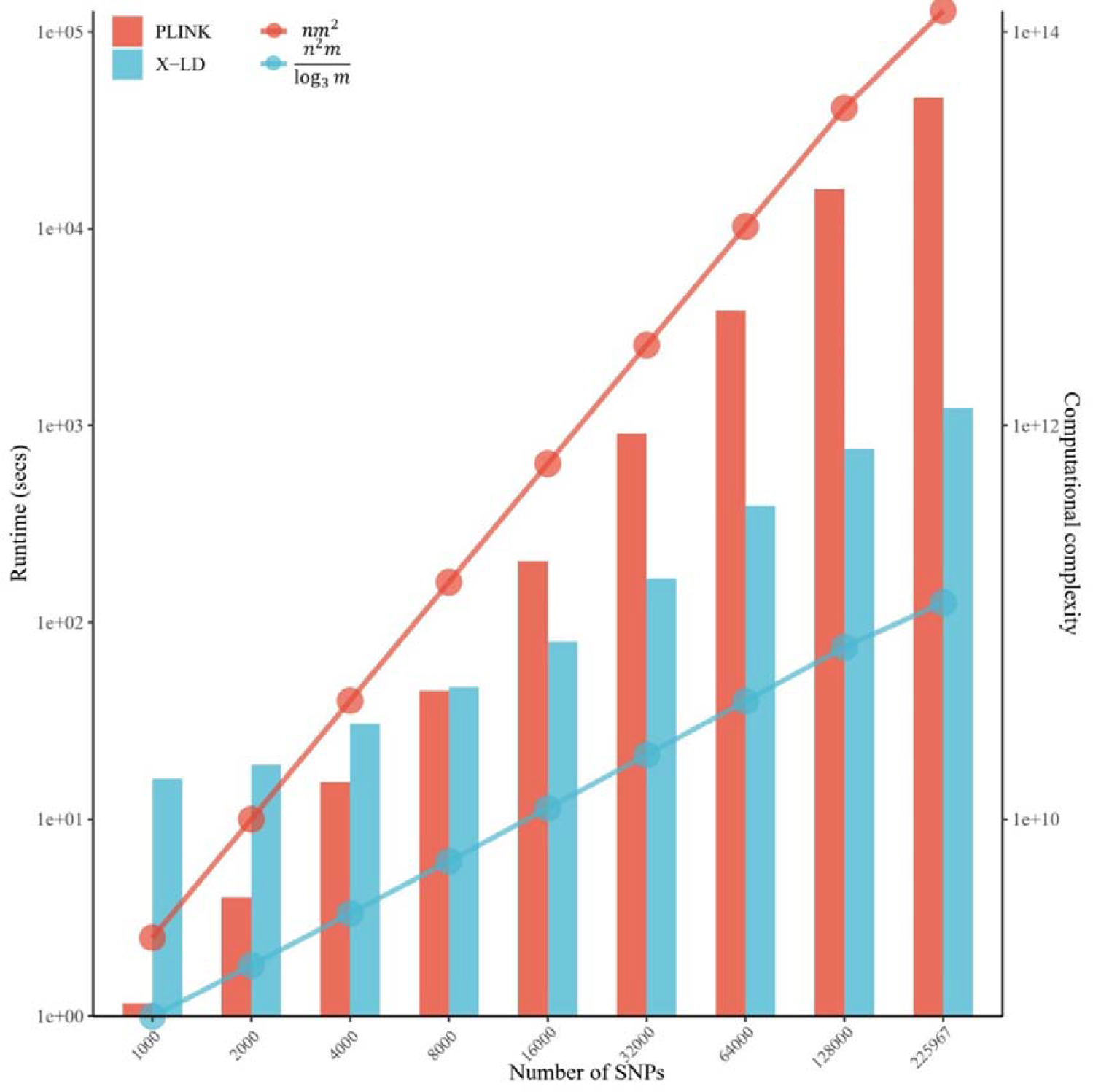
The computational efficiency of X-LD algorithm. Considering the high computational cost of PLINK, only the first chromosome was chosen. In the process of evaluating computational efficiency, we kept adding SNPs until the inclusion of entire chromosome. The bar chart and line chart show the actual calculation time and theoretical calculation complexity, respectively.

**Figure S3.**
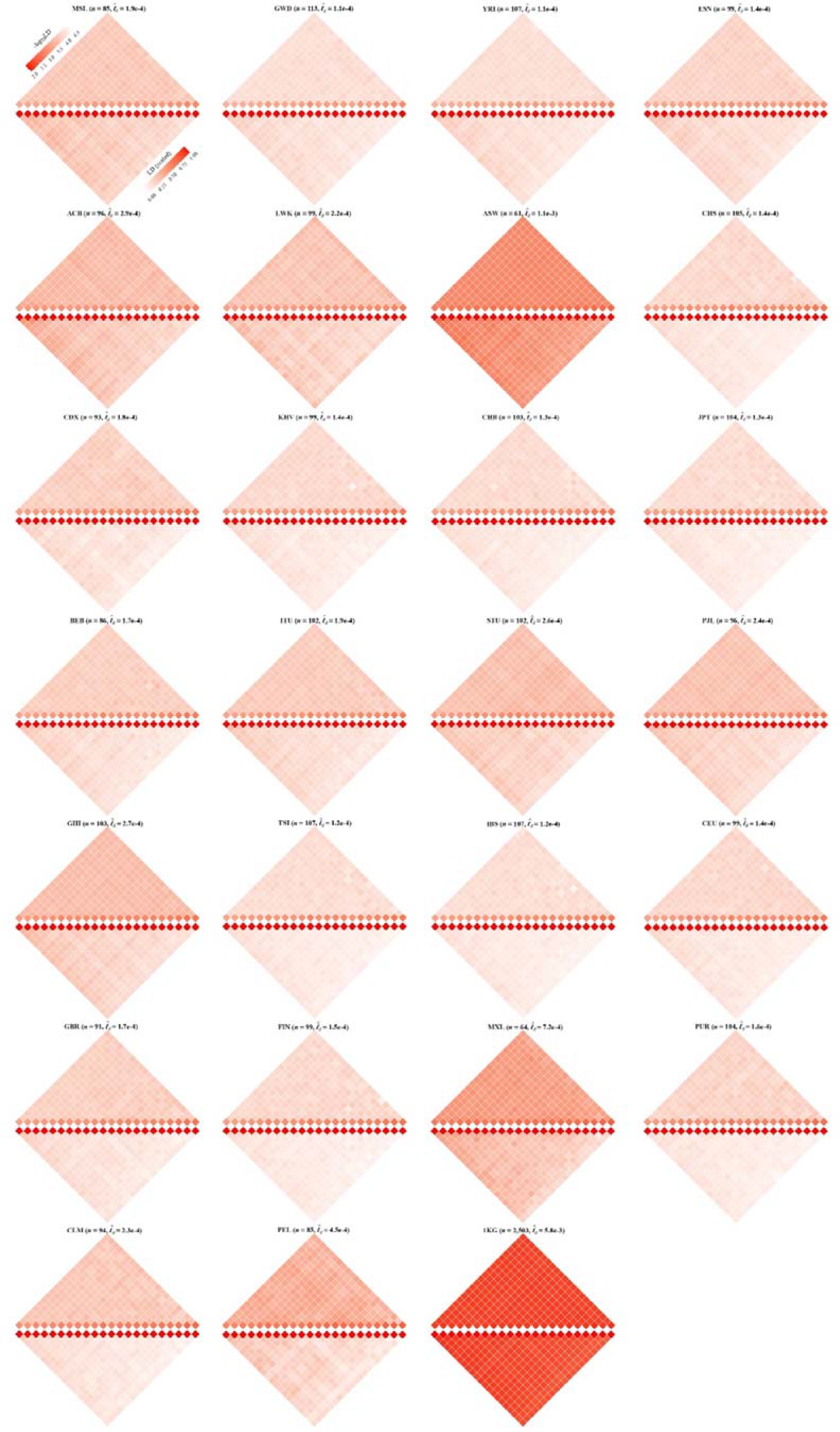
Chromosomal scale LD components for 26 cohorts in 1KG. The upper and lower parts of each figure represent the LD before and after scaling according to **Eq 8**. 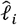 and 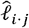 are represented by the diagonal and the off-diagonal elements, respectively. For visualization purposes, LD before scaling is transformed to a -log10-scale, with smaller values (red hues) representing larger LD, and a value of 0 representing that all SNPs are in LD.

**Figure S4.**
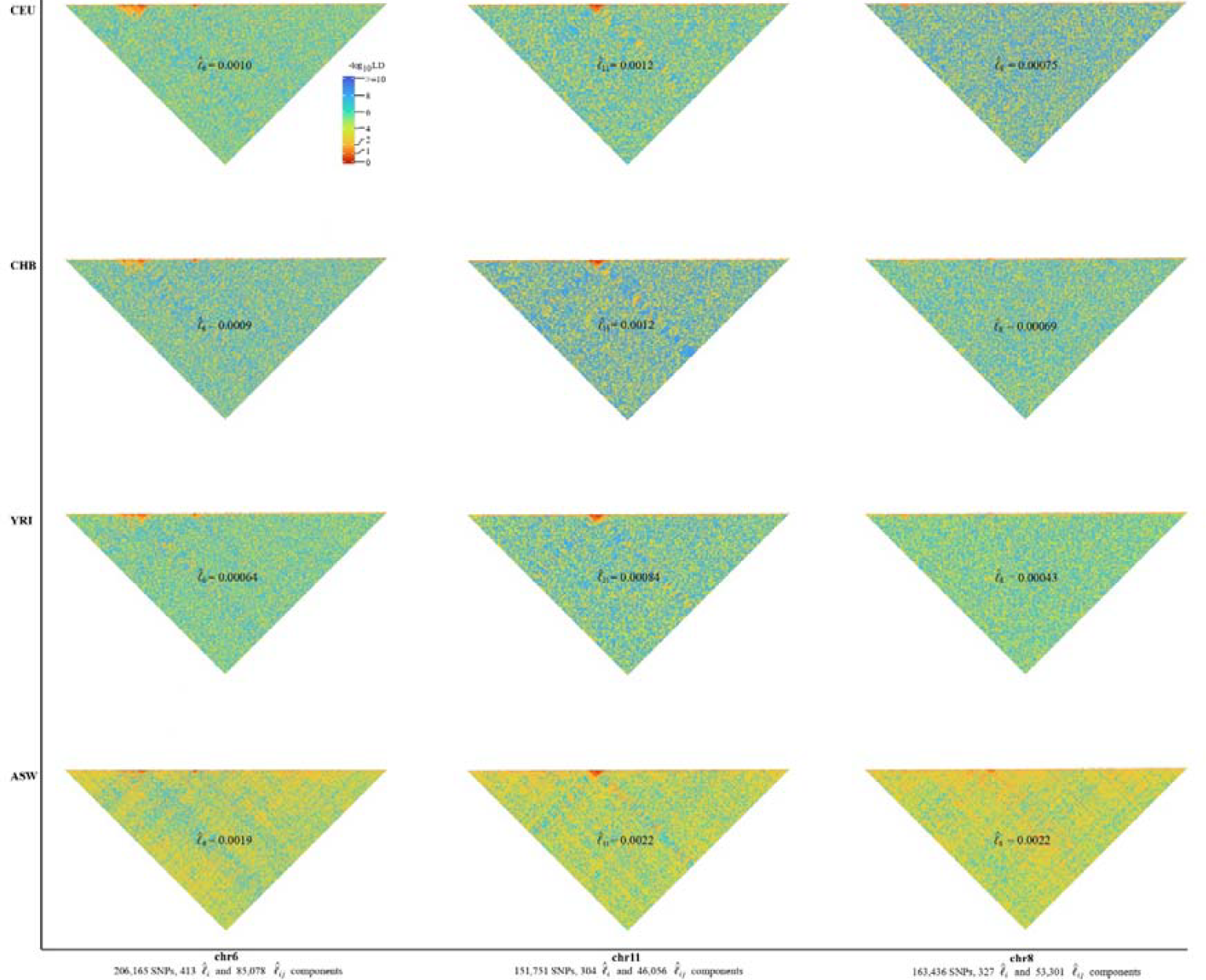
High-resolution illustration for LD grids for CEU, CHB, YRI, and ASW (m = 500). For each cohort, we partitioned each chromosome into consecutive LD grids (each LD grid containing 500 SNPs). For visualization purposes, LD is transformed to a -log10-scale, with smaller values (red hues) representing larger LD, and a value of 0 representing that all SNPs are in LD.

**Figure S5.**
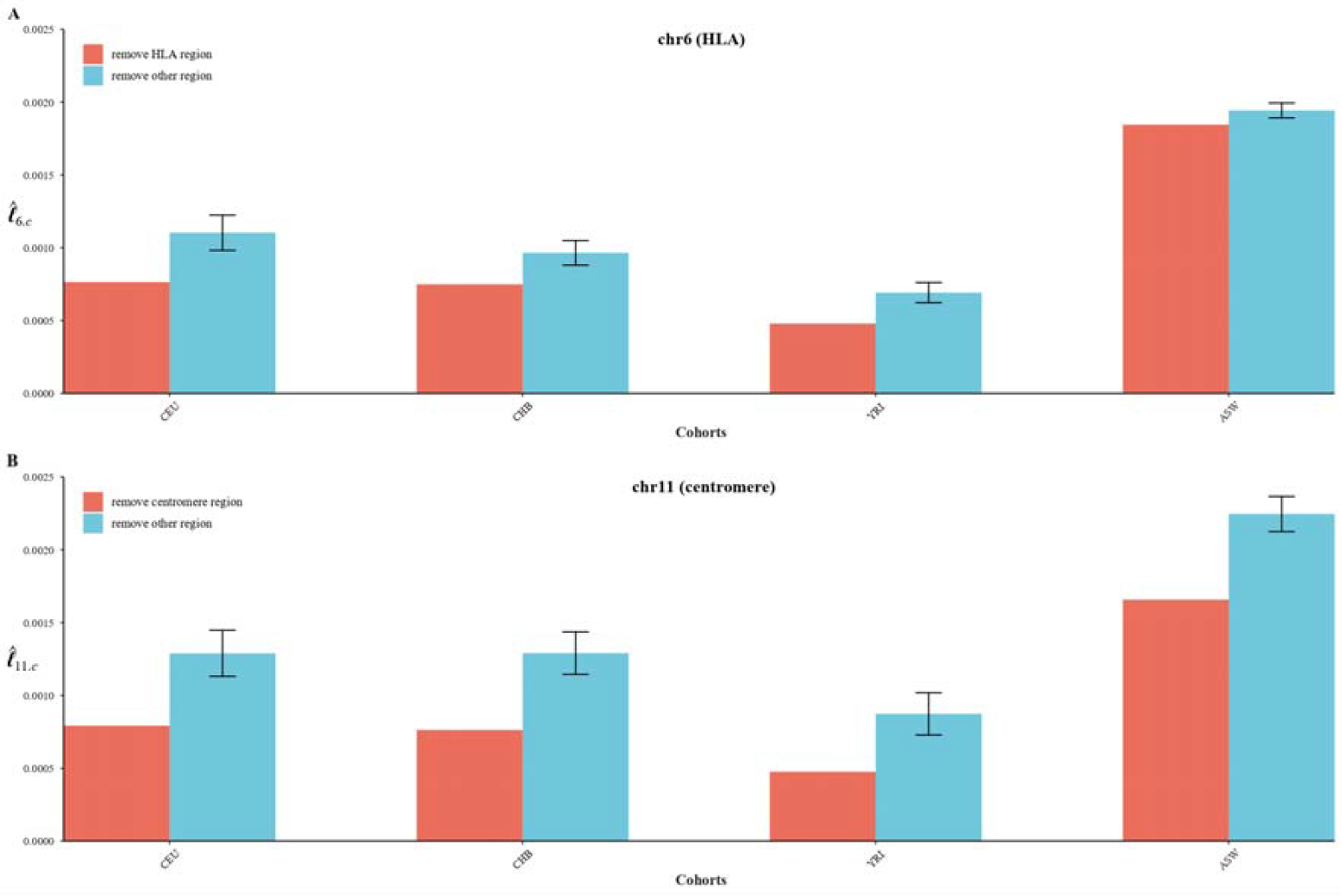
Influence of HLA region on chromosome 6 and centromere region on chromosome 11 on chromosomal LD in CEU, CHB, YRI, and ASW. When other region was removed, to avoid chance, the same number of consecutive SNPs as HLA region or centromere region were randomly removed from the genomic region, and this operation was repeated 100 times.

**Figure S6.**
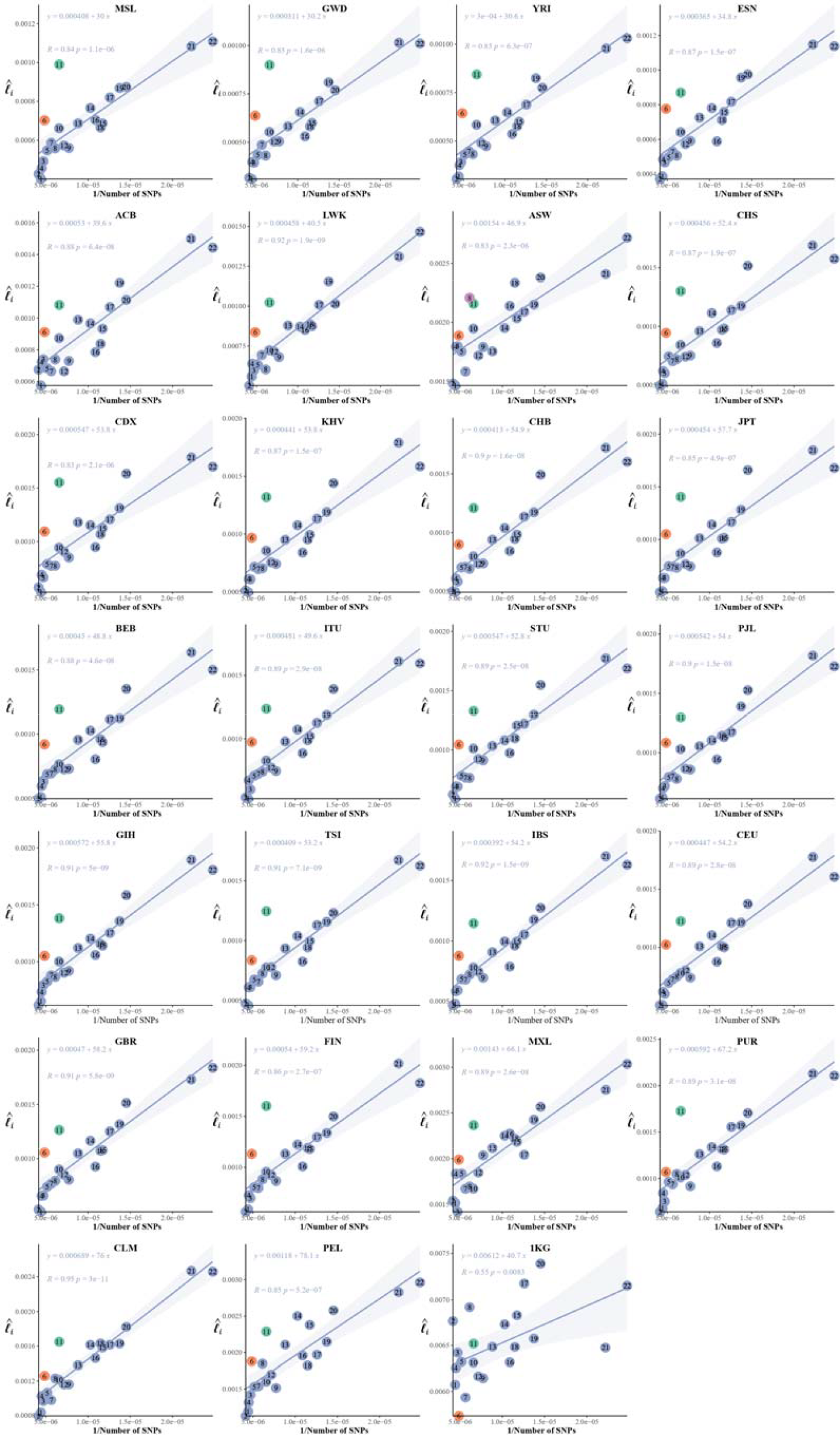
The correlation between the inverse of the SNP number and chromosomal LD in 26 cohorts of 1KG.

**Figure S7.**
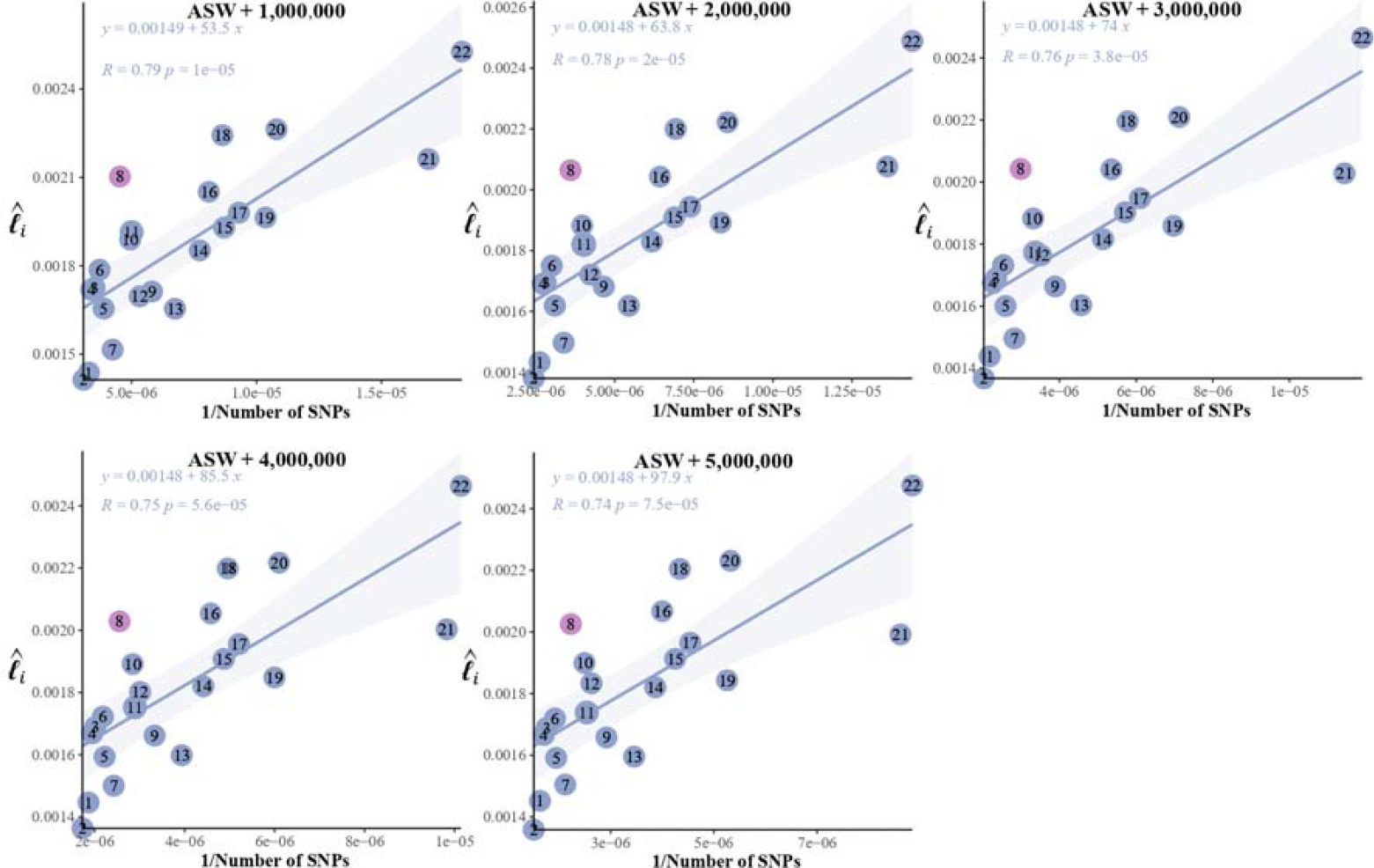
Influence of expanding of SNP numbers on the correlation between the inverse of the SNP number and chromosomal LD in ASW. Randomly selected SNPs that were presented in ASW but were not 2,997,635 consensus SNPs were added to the ASW cohort to demonstrate the stable pattern of chromosome 8.

