## Supplementary material for "Efficient estimation for large-scale linkage disequilibrium patterns of the human genome": R code for Figure 6C: LD_decay_regression.R

```
setwd("./")
library(ggplot2)
library(stringr)
library(RColorBrewer)
library(plotly)
library(magrittr)
library(ggpubr)
library(ggrepel)
library(ggpmisc)
#######################################################################
List_Files <- c("CEU","CHB","YRI","ASW")

# 1.Number of SNPs vs Intra-c LD
j=1
for(i in 1:length(List_Files)){
  Intra_C <- data.frame(matrix(NA,ncol = 4))
  colnames(Intra_C) <- c("snp","rsq","datasets","chr")
  Data <- read.table(paste0(List_Files[i],".2.maf.geno.xld"),header = T, fill = TRUE)[,c(1:5)]
  Data <- Data[Data[,1] == Data[,3],]
  Len = nrow(Data)
  Intra_C[1:Len,1] <- 1/Data[,2]
  Intra_C[1:Len,2] <- Data[,5]
  Intra_C[1:Len,4] <- Data[,1]
  Intra_C[1:Len,3] <- "background"
  
  # chr6 
  Key <- data.frame(matrix(NA,ncol = 4))
  colnames(Key) <- c("snp","rsq","datasets","chr")
  Key[1,] <- Intra_C[which(Intra_C[,4]==6),] 
  Key[1,3] <- "type1"
  # chr11
  Key[2,] <- Intra_C[which(Intra_C[,4]==11),] 
  Key[2,3] <- "type2"
  # chr8 (ASW)
  if(i == 4){
    Key[3,] <- Intra_C[which(Intra_C[,4]==8),] 
    Key[3,3] <- "type3"
  }
  cor = round(cor(Intra_C$snp,Intra_C$rsq),2)
  print(cor)
  p<-ggplot(data = Intra_C, mapping = aes(x = snp, y = rsq, colour = datasets,fill = datasets,label = chr)) + geom_point(size = 6) +
    geom_text(size = 4,family = "serif" ,color = "black", fill = "white") +
    scale_color_manual(values = c("#8DA1CB","#FD8D62","#66C3A5","#C98DC6")) +
    scale_fill_manual(values = c("#8DA1CB","#FD8D62","#66C3A5","#C98DC6")) +
    theme_classic() +
    geom_smooth(method=lm,formula = y ~ x, color="#8DA1CB",se=T,alpha = 0.1) + 
    stat_poly_eq(formula = y ~ x,
                 label.x = "left",
                 family="serif",
                 aes(label = paste(..eq.label..))) +
    #aes(label = paste(..eq.label..,"italic(R)~~`=`~",cor,..p.value.label..,sep = "~~")),
    #parse = TRUE) + 
    stat_cor(family="serif",method = "pearson",label.sep = " ",label.x.npc = 0.05,label.y.npc = 0.95) + 
    scale_y_continuous(expand = c(0,0)) +
    scale_x_continuous(expand = c(0,0)) +
    labs(x="1/Number of SNPs",y= "Intra-C LD",title=strsplit(List_Files[i],".2.maf.geno.adj.xld")[[1]][1])+
    theme(text=element_text(size = 12,family = "serif"),
          plot.title = element_text(family = "serif", face = "bold", vjust = -5,hjust = 0.5),
          legend.title = element_blank(),
          legend.position = "none") +
    theme(
      plot.margin = margin(t = 0,       
                           r = 0,       
                           b = 0,       
                           l = 0,       
                           unit = "cm")) + 
    geom_point(data = Key, mapping = aes(x = snp, y= rsq, colour = datasets, fill = datasets, label = chr),size = 6) +
    geom_text(family = "serif" ,color = "black", fill = "white") 
  
  #print(cor(Intra_C[,1],Intra_C[,2]))
  j = j + 1
  ggsave(paste(List_Files[i],".pdf"), plot = p)
}
```
