## Supplementary figures and images for "Efficient estimation for large-scale linkage disequilibrium patterns of the human genome"

### ASW .pdf

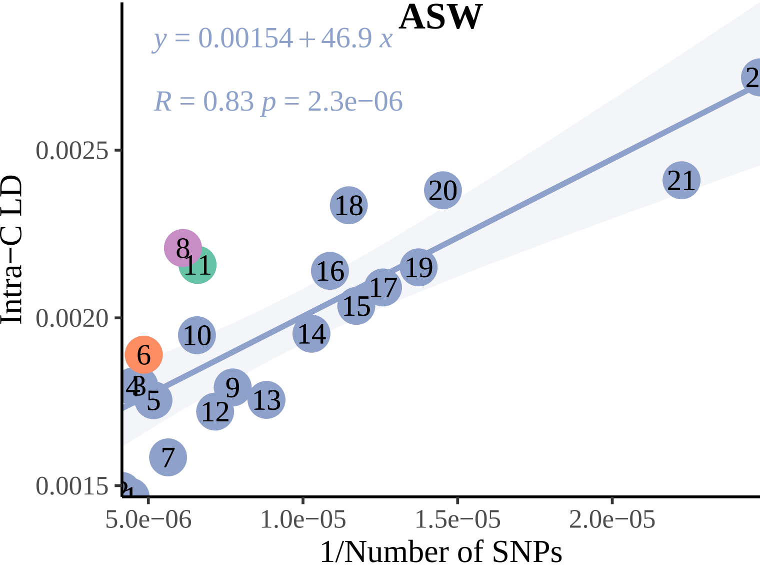

### CEU .pdf

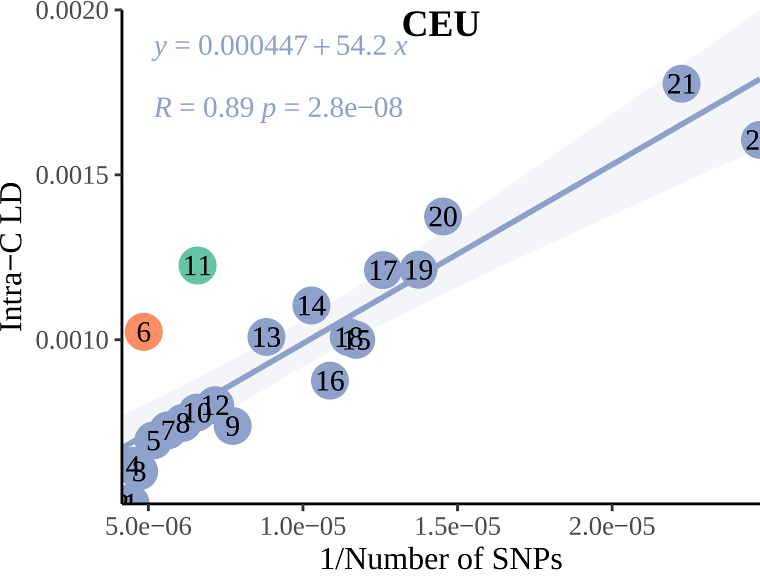

### CHB .pdf

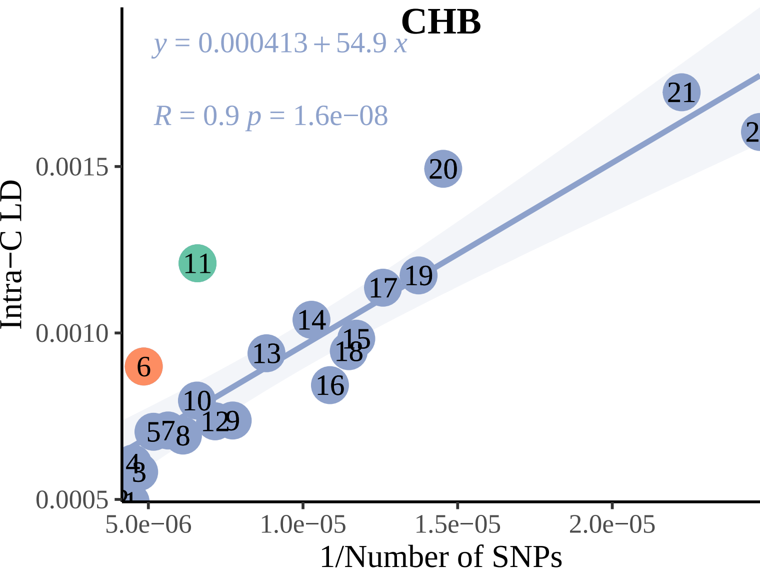

### YRI .pdf

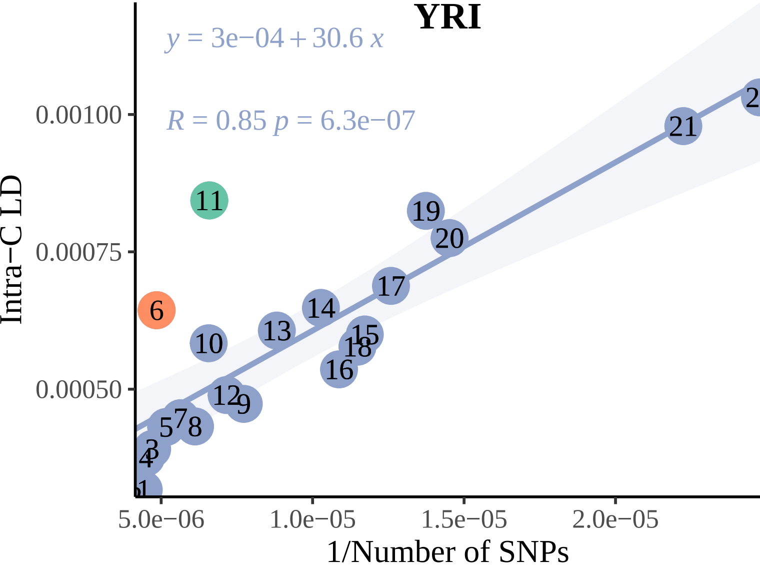
